# The Neurofibrillary Tangle Maturity Scale: A Novel Framework for Tangle Pathology Evaluation in Alzheimer’s Disease

**DOI:** 10.1101/2025.06.02.657435

**Authors:** Christina M. Moloney, Matthew H. Rutledge, Sydney A. Labuzan, Zhongwei Peng, Jessica F. Tranovich, Ashley C. Wood, Darren M. Rothberg, Ranjan Duara, Christian Lachner, Neill R. Graff-Radford, Dennis W. Dickson, Nicholas M. Kanaan, Rickey E. Carter, Melissa E. Murray

## Abstract

Neurofibrillary tangles are dynamic neuropathologic hallmarks of Alzheimer’s disease with a hypothesized lifespan morphologically-defined by three maturity levels: pretangles, mature tangles, and ghost tangles. To better understand the progression of tangle pathophysiology, we characterized tangle maturity level predilection of 15 tau antibodies recognizing a broad range of linear, phosphorylation, conformational, and truncation epitopes in the hippocampus of 24 postmortem brains. We developed the tangle maturity scoring system to semi-quantitatively evaluate each tangle maturity level. Based on proportions of tangle maturity levels, we classified antibodies as “early” (mostly pretangles and mature tangles), “middling” (mature tangles with pretangles and ghost tangles), and “advanced” (mostly ghost tangles and mature tangles) tangle maturity markers. To summarize tangle maturity predilection, we developed the tangle maturity scale to integrate individual tangle maturity scores. Correlations showed stronger relationships between tangle maturity scale and subsector thickness for more advanced tangle maturity markers in CA1 and subiculum, whereas Braak tangle stage remained consistently correlated throughout markers of the tangle lifespan. To aid in scoring hippocampi, we used machine learning to recognize tangle maturity levels, which performed comparably to a domain expert and showed similar relationships by Spearman correlation. Pattern recognition software was used to assess tangle and neuritic tau burden separately, which generally correlated with Braak stage and neuronal counts. However, tangle-derived tau burden more consistently correlated with hippocampal subsector thickness. In conclusion, we developed manual and automated scoring systems to evaluate tangle maturity levels, demonstrating early 4R, phosphorylated, and oligomeric tau accumulation preceding more advanced 3R and truncated tau. Our study provides supportive evidence of disease-relevant ordering of tau posttranslational modifications in the brain, which may have implications for theragnostic development. These findings underscore the promise of computerized quantitative analyses (i.e., pathomics) for high-throughput feature extraction from whole-slide images to enhance our understanding of microscopically observed morphologic changes.

## Introduction

Neurofibrillary tangles are composed of abnormal posttranslationally modified tau and are one of the neuropathologic hallmarks of Alzheimer’s disease [73]. Accumulation of tangles is associated with cognitive symptom onset and neuronal death [27, 30, 52, 56]. Tangle-bearing neurons are hypothesized to progress through three morphologically-defined maturity levels: pretangles, mature tangles, and ghost tangles [51]. Pretangles have diffuse or granular cytoplasmic tau staining, often accompanied by perinuclear tau accumulation [8]. Mature tangles are composed of densely packed bundles of tau fibers with nuclear shrinking or localization towards the cell membrane [1-3]. Ghost tangles are extracellular remnants of tangle-bearing neurons, appearing as loosely arranged bundles of fibers without a nucleus. Additionally, there are transitional intermediaries described between the tangle maturity levels [51]. Fluid and neuroimaging tau biomarkers are found to differentially associate with early and advanced markers of tangle maturity [39, 45, 47, 50, 51, 53, 54, 59], prompting further investigation into morphologic characteristics of posttranslationally modified tau. Previous studies have characterized the recognition of tangle maturity levels through strategic selection of tau antibodies [5, 7, 17, 33, 35, 71], as well as demonstrated region-specific differences underlying clinical heterogeneity in atypical forms of Alzheimer’s disease [6, 19, 43].

With numerous tau antibodies available that recognize subsets of tangle maturity levels [10, 17, 33, 35, 38, 51], we were motivated to characterize a wide range of tau antibodies in the context of hippocampal tissue health across Braak stages [12, 13]. With high potential for translational impact, we sought to develop methods for quantifying and summarizing tangle maturity levels and differentiating neuritic tau burden from tangle burden. As machine learning approaches coupled with robust descriptive observations continues to show promise in transformative neuropathology studies [48, 55, 62, 66, 72], we were also inspired to investigate computational quantitative methods (i.e., pathomics) to assess tau antibody characteristics. Pathomics is a burgeoning cross-disciplinary field that integrates a comprehensive approach involving the extraction, analysis, and characterization of vast amounts of quantitative data derived from high-resolution tissue images using artificial intelligence (AI) [14, 15, 22, 34, 37, 63, 68]. Thus, our overall goal was to develop quantification methods for tangle maturity levels and provide recommendations of tangle maturity markers to fully capture the tangle lifespan. To accomplish this, our first aim was to develop a manual tangle maturity scoring system to semi-quantify tangle maturity levels. We next aimed to develop the tangle maturity scale to quantify how advanced tangle maturity is in tissue. Finally, we introduced a machine learning modeling approach to automate the tangle maturity scoring and scale.

## Materials and Methods

### Series selection

Cases were selected as previously described [50]. Briefly, The FLorida Autopsied Multi-Ethnic (FLAME) cohort [64], as of May 27, 2020, was queried to identify 4 cases (2 females, 2 males) from each Braak tangle stage (I-VI). Cases with significant non-AD neurodegenerative pathology and non-AD tauopathies were excluded. Cases with age-related tauopathies were not excluded. This study series included 24 cases, as summarized in **Supplementary Table 1**.

### Tissue sectioning and immunohistochemistry

Tissue sectioning was performed as previously described [50]. Briefly, 5 µm formalin-fixed paraffin-embedded sections of the posterior hippocampus were deparaffinized and rehydrated using standard methods. Immunohistochemistry was performed on serial sections using the Lab Vision Autostainer 480S (Thermo Scientific). Specifications and vendor information for each antibody are reported in **Table 1** with antibody map shown in **Figure 1**. As tau antibodies rarely stain every tangle through ghost tangles, we attempted to stain all tangles with dual-labelling RD3 and RD4 together (RD3&RD4). Sections were developed (Biocare Medical, catalog # M3M530L [mouse] or M3R531L [rabbit]), counterstained with hematoxylin, and coverslipped using Cytoseal. Slides were cured a minimum of 48 hours at room temperature before scanning. To compare antibodies to an advanced tangle maturity marker used routinely in the brain bank, we also stained sections with 1% thioflavin-S, and TO-PRO-3, a nuclear stain, as previously described [50].

**Table 1.** Antibody information.

| Antibody | Epitope | Source Catalog # | RRID | Species | Clonality | Isotype | Dilution | Antigen Retrieval | Reference |
| --- | --- | --- | --- | --- | --- | --- | --- | --- | --- |
| 2E9 | aa 347-366 | Novus Biologicals NBP2-25162 | AB_2895250 | Mouse | Monoclonal | IgG1 | 1:100,000 | Formic Acid |  |
| AT180 | pT231 | ThermoFisher Scientific MN1040 | AB_223649 | Mouse | Monoclonal | IgG1κ | 1:5,000 | H<sub>2</sub>O steam | [29] |
| AT270 | pT181 | ThermoFisher Scientific MN1050 | AB_223651 | Mouse | Monoclonal | IgG1κ | 1:10,000 | H<sub>2</sub>O steam | [29] |
| AT8 | pS202, pT205 | ThermoFisher Scientific MN1020 | AB_223647 | Mouse | Monoclonal | IgG1k | 1:2,500 | H<sub>2</sub>O steam | [49] |
| GT-38 | Conformational | Abcam ab246808 | AB_2864300 | Mouse | Monoclonal | IgG1 | 1:20,000 | H<sub>2</sub>O steam | [28] |
| MC1 | Conformational aa 7-9, 312-322 | Benjamin Wolozin | AB_2314773 | Mouse | Monoclonal | IgG1 | 1:250 | H<sub>2</sub>O steam | [40] |
| MN423 | Truncation, aa 387-391 | Michal Novac |  | Mouse | Monoclonal | IgG1 | 1:500 | H<sub>2</sub>O steam | [42, 57, 58, 76] |
| PHF-1 | pS396, pS404 | Peter Davies | AB_2315150 | Mouse | Monoclonal | IgG1 | 1:1,000 | H<sub>2</sub>O steam | [32] |
| pT205 | pT205 | ThermoFisher Scientific 44-738G | AB_2533738 | Mouse | Polyclonal | IgG | 1:2,500 | H<sub>2</sub>O steam |  |
| pT217 | pT217 | ThermoFisher Scientific 44-744 | AB_2533741 | Mouse | Polyclonal | IgG | 1:500 | H<sub>2</sub>O steam |  |
| RD3 | aa 267-274, 306-316 continuous | Millipore Sigma 05-803 | AB_310013 | Mouse | Monoclonal | IgG | 1:5,000 | Formic Acid | [21] |
| RD4 | aa 275-291 | Millipore Sigma 05-804 | AB_310014 | Mouse | Monoclonal | IgG | 1:5,000 | Formic Acid | [21] |
| Tau-66 | Conformational, aa 155-244, 305-314 | Nicholas Kannan | AB_2832938 | Mouse | Monoclonal | IgM | 1:2,500 | None | [26] |
| TauC3 | Truncation, aa 412-421 | Nicholas Kannan | AB_2832930 | Mouse | Monoclonal | IgG1 | 1:5,000 | H<sub>2</sub>O steam | [24] |
| TOC1 | aa 209-224 | Nicholas Kannan | AB_2832939 | Mouse | Monoclonal | IgG | 1:2,500 | H<sub>2</sub>O steam | [61, 74] |

Abbreviations: aa, amino acid; Ig, immunoglobulin; pT, phosphorylated threonine; pS, phosphorylated serine; RRID, Research Resource Identifier

**Fig. 1.**
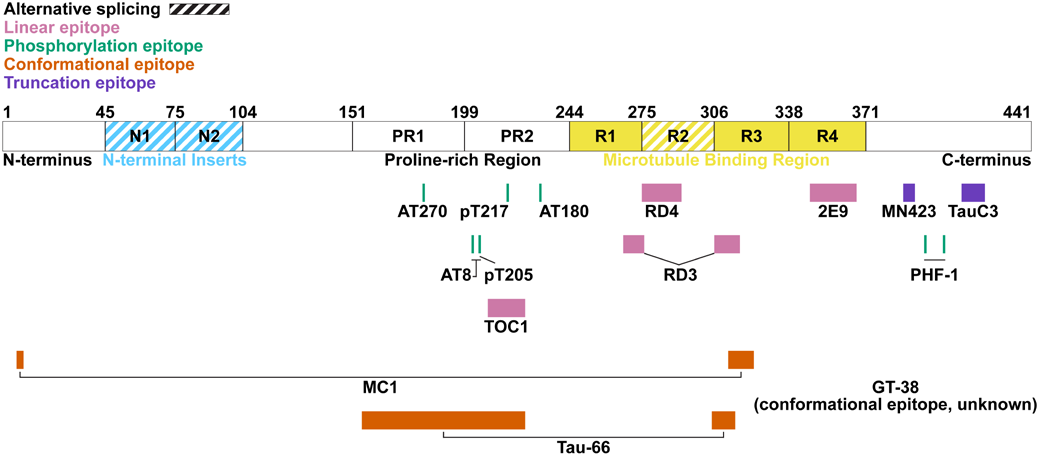
Antibody epitopes plotted along the linear 2N4R tau 441 amino acid structure with alternatively spliced regions indicated by diagonal lines. Light blue represents the amino-terminal (N-terminal) inserts, yellow the microtubule binding repeat region, pink the linear epitopes, green the phosphorylation epitopes, orange the conformational epitopes, and purple the truncation epitopes. Epitopes are detailed in **Table 1**. Figure adapted from [51]. Note: TOC1 recognizes a linear epitope upon conformation following N. terminal cleavage event [74]. Abbreviations: N, amino; C, carboxy, PR, proline-rich region; R, repeat domain

### Digital pathology

Hematoxylin and eosin (H&E) and immunohistochemically stained slides were digitally scanned at 20x magnification with the Aperio AT2 scanner (Leica Biosystems). The CA1, CA2/3, CA4, and subiculum were annotated using the pen tool, excluding artifacts with the negative pen tool in ImageScope (Leica Biosystems, version 12.4.6.7001). Additionally, the thickness of the strait of the CA1 and subiculum was determined by averaging 3 measurements perpendicular to the strait of each region, extending to the gray-white matter junctions. Brightfield scans were batch analyzed using eSlideManager (Leica Biosystems). Burden was measured by analyzing digitized slides with a positive pixel count (v9) or color deconvolution (v9) macro, custom designed for each antibody to recognize the 3, 3’-diaminobenzidine staining (**Supplementary Table 2, 3**). Pattern recognition software used to differentially measure tau burden in tangles and neuritic pathology is further described in “Pathomics” section below.

**Table 2.** Neurofibrillary tangle maturity semi-quantitative scores.

| Interpretation | Score | # Tangles/1.62 $\text{mm}^2$ | |
| --- | --- | --- | --- |
|  |  | Manual | Model |
| None | 0 | 0 | 0-0.12 |
| Rare | 0+ | 0-1 | 0.12-1.19 |
| Sparse | + | 1-4 | 1.19-5.78 |
| Moderate | ++ | 4-9 | 5.78-15.13 |
| Frequent | +++ | $\geq 10$ | $\geq 15.13$ |

**Table 3.** Neurofibrillary tangle maturity scale.

| Pretangles | Mature Tangles | Ghost Tangles |  |  |  |
| --- | --- | --- | --- | --- | --- |
|  |  | 0/0+ | + | ++ | +++ |
| 0/0+ | 0/0+ | 0/1 | 6 | 6 | 6 |
| + | 0/0+ | 2 | 5 | 6 | 6 |
| ++ | 0/0+ | 2 | 2 | 5 | 5 |
| +++ | 0/0+ | 2 | 2 | 5 | 5 |
| 0/0+ | + | 4 | 5 | 5 | 6 |
| + | + | 3 | 5 | 5 | 6 |
| ++ | + | 3 | 3 | 5 | 5 |
| +++ | + | 2 | 2 | 5 | 5 |
| 0/0+ | ++ | 4 | 5 | 5 | 5 |
| + | ++ | 3 | 4 | 5 | 5 |
| ++ | ++ | 3 | 3 | 5 | 5 |
| +++ | ++ | 3 | 3 | 5 | 5 |
| 0/0+ | +++ | 4 | 4 | 5 | 5 |
| + | +++ | 4 | 4 | 5 | 5 |
| ++ | +++ | 3 | 4 | 5 | 5 |
| +++ | +++ | 3 | 3 | 5 | 5 |

Tangle maturity levels were scored as detailed in **Table 2**.

Fluorescent-stained sections were digitally scanned at 20x magnification using the Pannoramic 250 Flash III (3DHistech) (for scanning parameters, see [50]). Annotations for CA1, CA2/3, CA4, and subiculum were made with the closed polygon tool for the thioflavin-S/TO-PRO-3-stained sections using CaseViewer (3DHistech, version 2.3.0.99276).

### Neurofibrillary tangle maturity scores and scale

We used a semiquantitative approach to score the presence of each tangle maturity level. Samples were rated as sparse (0<1 tangle), mild (2-4 tangles), moderate (5-9 tangles) or severe (≥ 10 tangles) per 1.62 mm^2^ area (area obtained from a 10x view in ImageScope, **Table 2**, manual). Using the scores for pretangles, mature tangles, and ghost tangles, a composite scaled value was calculated (**Table 3**).

### Computerized quantitative analyses (Pathomics)

#### Tangle and neuritic burden pattern recognition using GENIE software

To analyze the burden of the tangle or neuritic pathology, we utilized GENIE [36], a pattern recognition software within Leica’s eSlideManager [20]. Examples of tangles, neuritic plaques, blood vessels, and neuropil from posterior hippocampi stained with AT8, PHF-1, and MN423 were annotated and trained for a total of 2,000 iterations. To quantify tangle pathology, the base macro was used with the addition of the tangle classifier, which included all tangle maturity levels. To quantify neuritic pathology, the base macro was used with the addition of the neuritic plaques and neuropil classifiers.

#### Neurofibrillary tangle maturity model (Tangle-mat AI) training methodology

We trained a fully-supervised object detection model to locate and classify tangles from whole slide images. To do this, we curated a set of tiles from whole slide images that were annotated by a domain expert with bounding boxes to delineate the location and type of tangle within each tile. We selected archived tau-stained slides from the Translational Neuropathology Laboratory database, and tiles from the posterior hippocampus were randomly chosen (**Fig. 2**). In total, there were 18 different stains in the training data and the majority of the slides were stained with 2E9 and AT8 antibodies (**Supplementary Table 4**). This was intentional to enhance the “Tangle-mat AI” model generalizability beyond this study as these two antibodies cover the tangle maturity

**Fig. 2.**
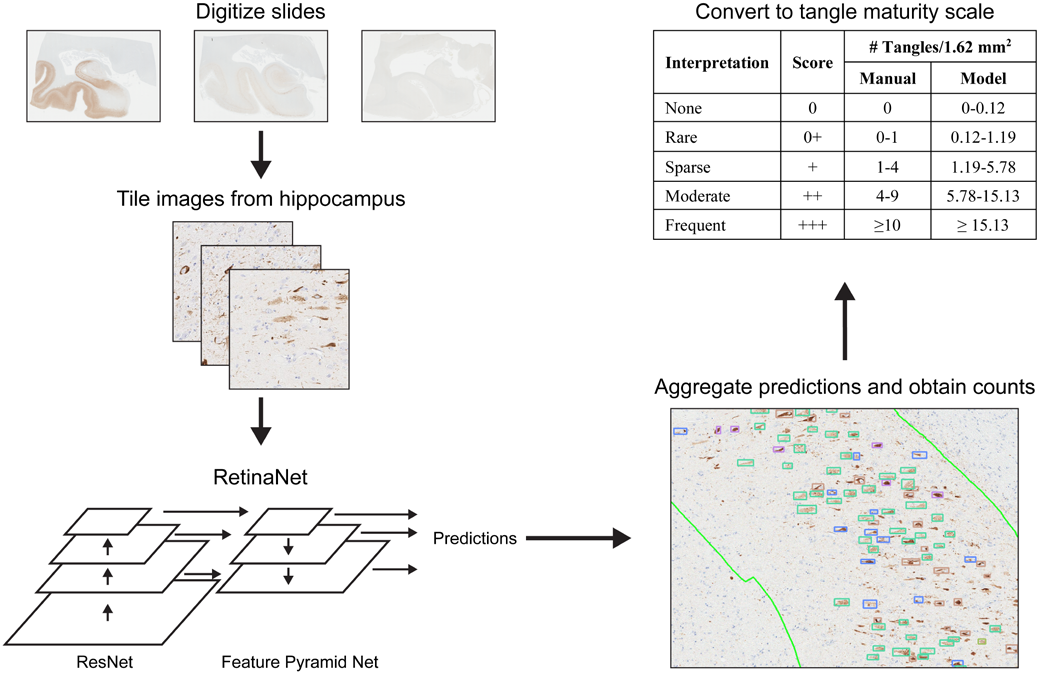
Pathomics workflow for processing whole slide images using the Tangle-mat AI model. Digitized whole slide images are divided into 600×600 pixel tiles, each tile is analyzed with a fine-tuned RetinaNet model, and predictions are aggregated to generate whole-slide counts. Counts were used by the Tangle-mat AI model to convert to tangle maturity scale, as shown in the table.

#### Acronyms: AI, artificial intelligence

lifespan. Additionally, tangle morphology across the tangle maturity levels is consistent across tau stains in our database, indicating that a model trained primarily on AT8 and 2E9 should generalize well to other stains. The training dataset for the Tangle-mat AI model consisted of 2,827 unique images from 714 unique slides. A total of 862 (30%) of these tiles came from slides included in this study, while the remaining 70% came from slides not included. The tiles were annotated with bounding boxes by a domain expert for three maturity levels and intermediaries (pretangle, intermediary 1, mature tangle, intermediary 2, and ghost tangle, **Supplementary Table 5**). Tangle maturity levels were identified as previously described [51]. Briefly, pretangles were labeled if the neuron had diffuse or granular staining of tau and may include perinuclear accumulation. Intermediary 1s had intensely stained aggregates but filled less than half the neuron, with a nucleus visible. Mature tangles were intensely stained and took up over 50% of the neuron, with a nucleus visible. Intermediary 2s were intensely stained through the neuron but lacked a nucleus. Ghost tangles were more faintly stained, had loosely arranged bundles of fibers, and lacked a nucleus. Lesions that resembled tangles but could not be confidently assigned a maturity level were labeled “unclassified.”

An initial 80/20 train-validation split was used to determine optimal learning rate and model architecture. The learning rate was determined using both a learning rate finder and grid search. All candidate models trained over 100 epochs. We evaluated all object detection models available in the TorchVision library and found the best performance with RetinaNet [44] with a ResNet50 backbone (retinanet_resnet50_fpn, finetuned from ‘COCO_V1’ weights — TorchVision 0.21 documentation). This is consistent with the literature showing focal loss used in RetinaNet aids with unbalanced classes, similar to the distribution of tangle maturities in the training data [44]. Upon determining an acceptable learning rate, data was partitioned using 5-fold cross validation. Data was partitioned such that images from a slide did not appear in both the training and validation set. We used a 5-fold cross validation to avoid overfitting the initial model selection and to quantify uncertainty in model performance metrics. A threshold of 0.25 was chosen to maximize model accuracy across all folds by filtering out low probability tangles (only tangles with a probability score greater than 0.25 were counted). All models were trained for the full 100 epochs using the selected learning rate.

### Finally, a full Tangle-mat AI model was fit with all data used for inference on whole slide images

#### Whole Slide Inference

For inference on whole slide images, regions of interest were divided into 600×600 pixel tiles with a 300-pixel stride. The full model trained on all the data was run on the tiles. Results from overlapping tiles were aggregated using non-maximum suppression to prevent double-counting tangles [67].

#### Model generated tangle maturity score and tangle maturity scale values

To convert tangle counts from the Tangle-mat AI model to the tangle maturity scale, we first converted the counts to the tangle maturity scores. To determine appropriate thresholds for class i vs class i+1 we used the geometric mean of the 80^th^ quantile of normalized Tangle-mat AI counts for class i and the 20^th^ quantile of class i+1. This was done using all slides/stains to avoid overfitting thresholds for each stain. This corresponds to the following cutoffs: none = 0-0.12 per 1.62 mm^2^; rare = 0.12-1.19 per 1.62 mm^2^; sparse = 1.19-5.78 per 1.62 mm^2^; moderate = 5.78-15.13 per 1.62 mm^2^; and severe = > 15.13 per 1.62 mm^2^ (**Table 2**, model). Based on this acceptable level of agreement, we performed Spearman correlations of the model’s tangle maturity scale for all antibodies, Braak stage, hippocampal subsector thickness, and neuronal count.

#### Neuron object detection model

To facilitate neuronal counts in hippocampal regions, we developed an object detection model to identify neurons in H&E-stained whole slide images. Tiles from whole slide images were randomly selected from archived slides and neurons were annotated with bounding boxes by a domain expert. The final dataset totaled 462 unique images from 56 unique whole slide images, totaling 3,839 labeled neurons. A training protocol similar to the tangle maturity object detection model was used. After tuning parameters on a single fold, 5-fold cross validation was used to assess model performance. The model trained to detect neurons in H&E-stained images had a recall of 86% and a precision of 72% at a threshold of 0.3 (only neurons with a probability score greater than 0.3 were counted) (**Supplementary Fig. 1**).

### Statistical analysis

Tangle maturity proportions were calculated by assigning a value to each score (none = 0, rare = 1, sparse = 2, moderate = 3, frequent = 4). For each antibody, the sum of scores for each tangle maturity level was divided by the total summed score and multiplied by 100. Spearman correlations were performed to correlate burden or tangle maturity scale for all 15 stains with Braak stage, subsector thickness, and neuronal counts. Correlations are described as negligible (0-0.1), weak (0.1-0.39), moderate (0.40-0.69), strong (0.7-0.89) and very strong (0.9-1) [65]. Statistical analyses were performed using R (version 4.4.1). P values <0.05 were considered statistically significant.

### Figure development

Figures were created using Adobe Illustrator 2025 (version 29.3.1). 10x and 20x view snapshots of brightfield scans were acquired using ImageScope (Leica Biosystems, version 12.4.3.5008). Plots were created using R (version 4.4.1).

## Results

### Antibody characterization

The 15 tau antibodies characterized included those that recognized linear epitopes (RD3, RD4, 2E9, TOC1 [upon conformational change]), phosphorylation epitopes (AT270 [pT181], pT205, pT217, AT180 [pT231], AT8 [pS202/pT205], PHF-1 [pS396/pS404]), conformational epitopes (MC1, Tau-66, GT-38), and truncation epitopes (TauC3, MN423) (**Fig. 1, Table 1**). Most antibodies stained all tangle maturity levels; however, one tangle maturity level was often rarely recognized compared to the others. pT181, pT205, pT231, AT8, TOC1, and MC1 did not stain ghost tangles in this case series (**Fig. 3**). Anecdotally, we observed rare tau aggregates of abnormal size with advanced tangle maturity markers (2E9, MN423) as exaggerated ghost tangles, informally referred to as “gigantaurs.” All antibodies recognized neuropil threads and neuritic plaques. All antibodies but 2E9, Tau-66, TauC3, and MN423 recognized tangle-associated neuritic clusters, and all but GT-38, Tau-66, and MN423 recognized tangle-bearing neurons with granulovacuolar degeneration bodies (**Fig. 4**). We additionally evaluated thioflavin-S, a fluorescent dye that binds β-pleated sheets and is used for Braak staging in the Mayo Clinic brain bank. Thioflavin-S stained mature tangles, ghost tangles, neuropil threads, neuritic plaques, and granulovacuolar degeneration bodies (**Supplementary Fig. 2**).

**Fig. 3.**
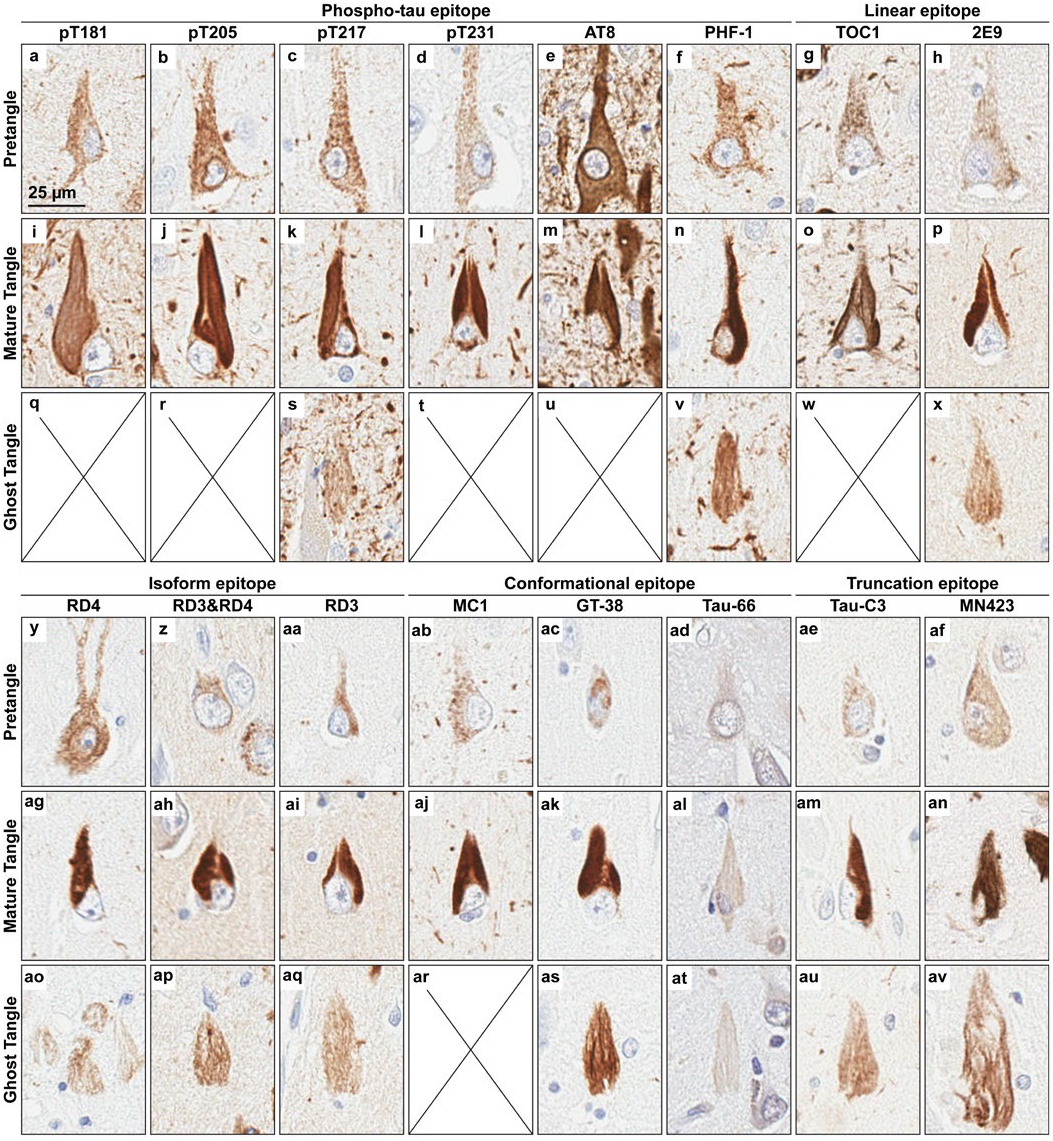
Tangle maturity level examples. Representative images from the series of pretangles **(a-h, y-af**), mature tangles (**i-p, ag-an**), and ghost tangles (**q-x, ao-av**) stained with pT181 (**a, i, q**), pT205 (**b, j, r**), pT217 (**c, k, s**), pT231 (**d, l, t**), AT8 (**e, m, u**), PHF-1 (**f, n, v**), TOC1 (**g, o, w**), 2E9 (**h, p, x**), RD4 (**y, ag, ao**), RD3&RD4 combined (**z, ah, ap**), RD3 (**aa, ai, aq**), MC1 (**ab, aj, ar**), GT-38 (**ac, ak, as**), Tau-66 (**ad, ail, at**), TauC3 (**ae, am, au**), and MN423 (**af, an, av**). Rare tau aggregates of abnormal size were observed with advanced tangle maturity markers as exaggerated ghost tangles, informally referred to as “gigantaurs” (**av**). Scale bar measures 25 µm and applies to all images.

**Fig. 4.**
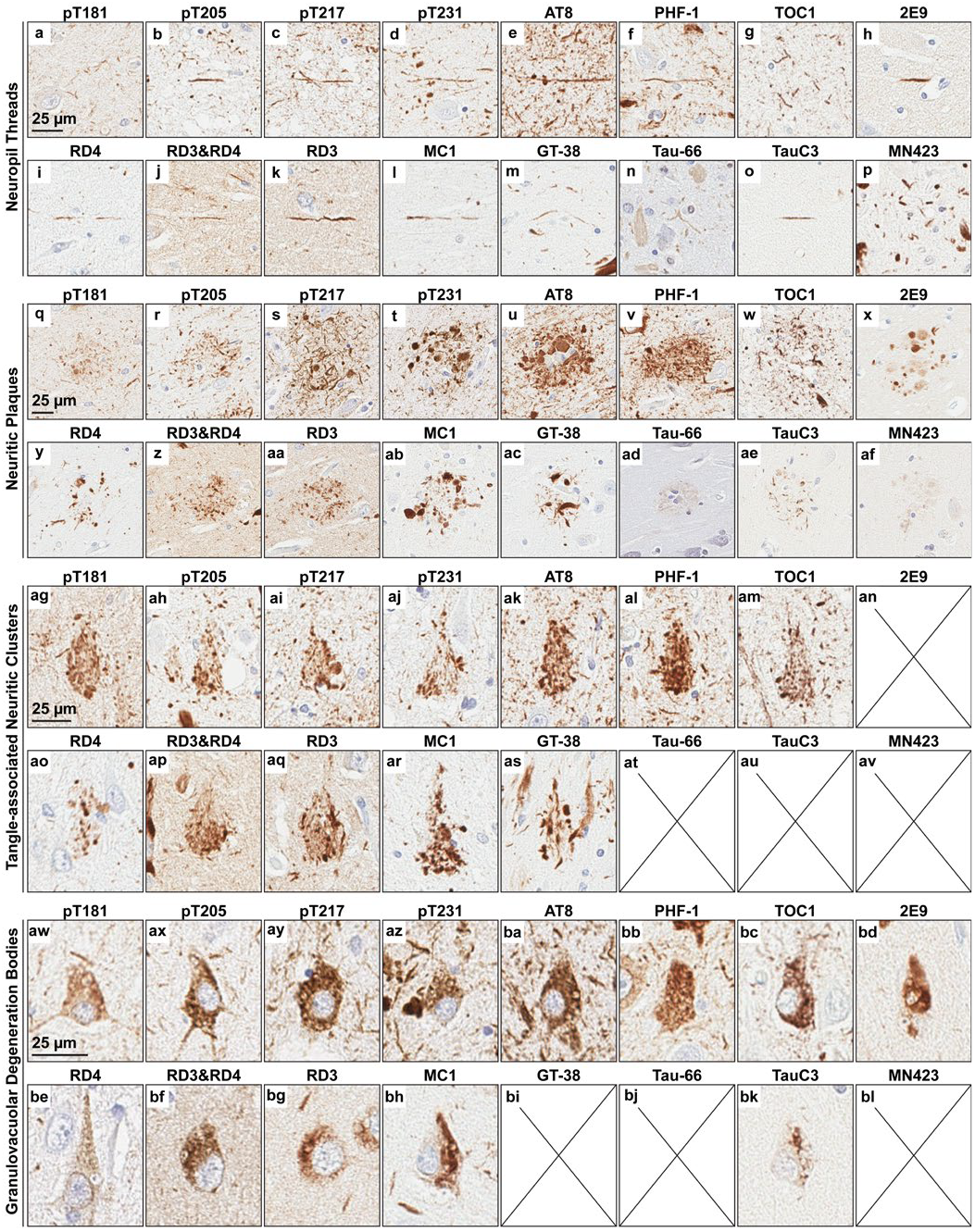
Non-tangle pathology examples. Representative images from the series of neuropil threads (**a-p**), neuritic plaques (**q-af**), tangle-associated neuritic clusters (**ag-av**), and tangles-bearing neurons with granulovacuolar degeneration (**aw-bl**) stained with pT181 (**a, q, ag, aw**), pT205 (**b, r, ah, ax**), pT217 (**c, s, ai, ay**), pT231 (**d, t, aj, az**), AT8 (**e, u, ak, ba**), PHF-1 (**f, v, al, bb**), TOC1 (**g, w, am, bc**), 2E9 (**h, x, an, bd**), RD4 (**i, y, ao, be**), RD3&RD4 combined (**j, z, ap, bf**), RD3 (**k, aa, ai, bg**), MC1 (**l, ab, ar, bh**), GT-38 (**m, ac, as, bi**), Tau-66 (**n, ad, at, bj**), TauC3 (**o, ae, au, bk**), and MN423 (**p, af, ab, bl**). Scale bars measures 25 µm and applies to all images within the specified pathology.

### Tangle maturity score and tau antibody ordering

We developed a 4-point scoring system to quantify each tangle maturity level and evaluated the CA1, CA2/3, CA4, and subiculum as these regions are differentially vulnerable throughout the course of Alzheimer’s disease [13]. We ordered the antibodies on increasing tangle maturity level recognition in the CA1 as it accumulates around half the number of tangles compared to the subiculum, allowing for a more dynamic range of tangle maturity assessment [50] (**Fig. 5a**). Antibodies with >50% pretangles were classified as “early” tangle maturity markers and included RD4, AT8, pT181, pT205, pT217, pT231, TOC1, and MC1 (**Fig. 6**).

**Fig. 5.**
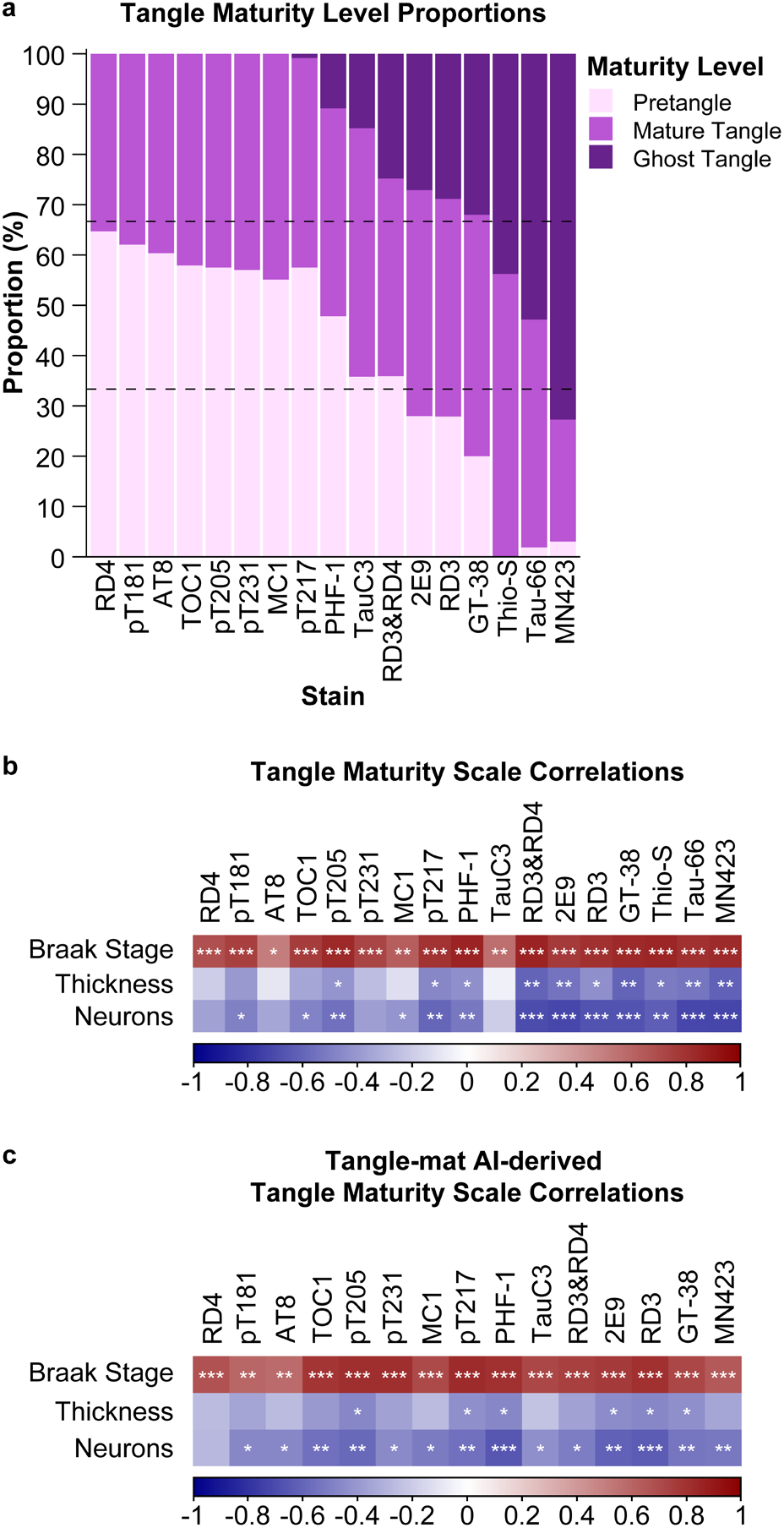
Tangle maturity quantification and correlations in the CA1. (**a**) Stacked bar graph of the proportion of pretangles, mature tangles, and ghost tangles for each stain. (**b**) Correlogram of Spearman correlations between the tangle maturity scale in the CA1 with Braak stage and tissue health (i.e., CA1 thickness, neurons). (**c**) Correlogram of Spearman correlations using the model derived tangle maturity scale. *, p<0.05; **, p<0.01; ***, p<0.001.

**Fig. 6.**
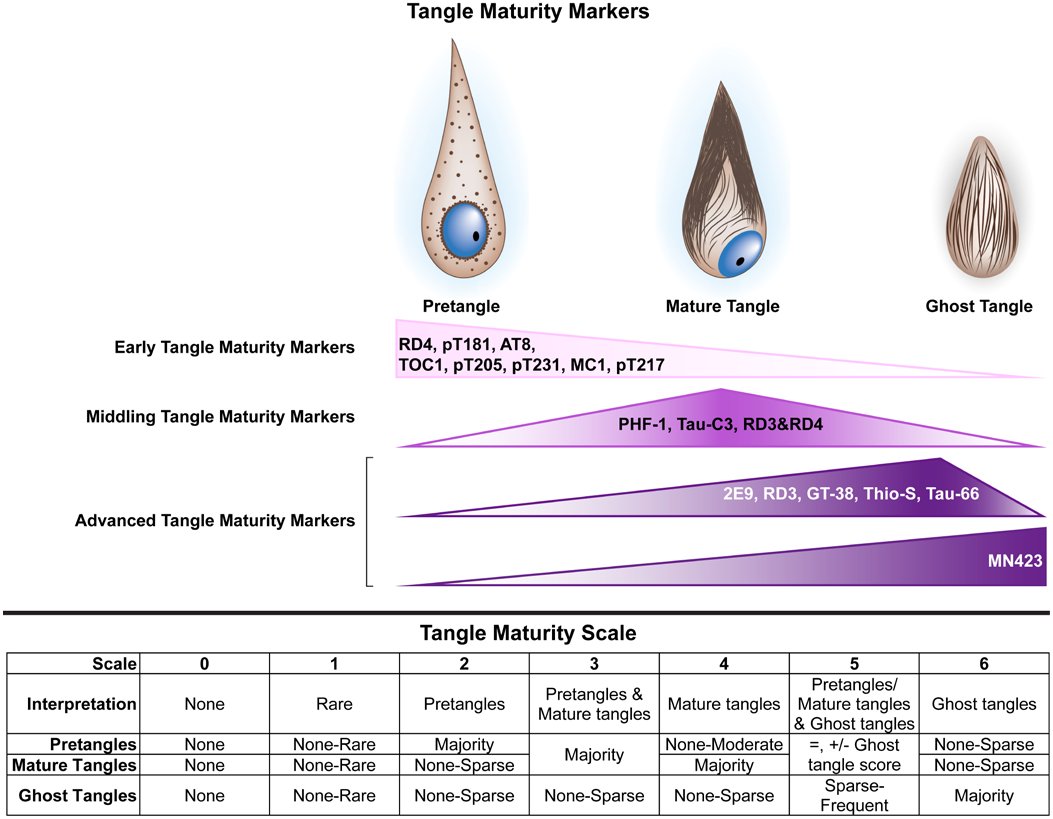
Tangle maturity summary. The tangle maturity markers are classified as “early”, “middling”, or “advanced” tangle maturity markers based on the tangle maturity proportions in the CA1 (**Fig. 5a**). Early tangle maturity markers include RD4, pT181, AT8, TOC1, pT205, pT231, MC1, and pT217. Middling tangle maturity markers include PHF-1. Tau-C3, and RD3&RD4. Advanced tangle maturity markers include 2E9, RD3, GT-38, thioflavin-S, Tau-66, and MN423. The tangle maturity scale is presented from 0-6, with 0 indicating no tangles, 1 indicating rare tangles, 2 indicating a majority of pretangles, 3 indicating a mix of pretangles and mature tangles, 4 indicating a majority of mature tangles, 5 indicating a mix of pretangles and mature tangles with ghost tangles, and 6 indicating a majority of ghost tangles. Figure adapted from [51]. Abbreviations: Thio-S, thioflavin-S

Antibodies with 50-66% mature and ghost tangles were classified as “middling” tangle maturity markers, and included PHF-1, TauC3, and dual-labeled RD3&RD4. Antibodies with >66% mature and ghost tangles combined were classified as “advanced” tangle maturity markers, and included 2E9, RD3, GT-38, Tau-66, and MN423. The other hippocampal subsectors followed a similar pattern, except RD4, pT205, and pT231 in the subiculum that recognized ghost tangles (**Supplementary Fig. 3a, 4a, 5a**).

### Tangle maturity scale

To provide a numeric summary value for the tangle maturity scores, we developed a novel 7-point scale by compositing the tangle maturity scores for pretangles, mature tangles, and ghost tangles (**Table 3, Fig. 6**). Correlations between the tangle maturity scale and Braak stage were moderate to strong in the CA1, moderate to very strong in the subiculum and CA4, and moderate in the CA2/3 (**Fig. 5, Supplementary Fig. 3b, 4b, 5b**). CA1 and subiculum thickness had moderate to strong inverse correlations with the tangle maturity scale (**Fig. 5, Supplementary Fig. 3b**), and these relationships tended to be stronger for antibodies targeting more advanced tangle maturity markers. The CA2/3 thickness had moderate to strong inverse correlations with antibodies spread through the tangle lifespan (pT181, pT231, PHF-1, Tau-66, **Supplementary Fig. 4b**). Neuronal counts in the CA1 were moderately to strongly correlated with all antibodies except RD4, AT8, and pT231, and the strength of correlation increased with more advancing tangle maturity markers (**Fig. 5b**). In the subiculum, neuronal counts were moderately correlated with middling and advanced tangle maturity markers, excluding TauC3 and GT-38 (**Supplementary Fig. 3b**). Neuronal counts in the CA2/3 were moderately correlated with few antibodies/stains across the tangle lifespan (pT231, TauC3, GT-38, thioflavin-S, **Supplementary Fig. 4b**). No significant correlations were observed between neuronal count and tangle maturity scale in the CA4 (**Supplementary Fig. 5b**).

#### Tangle-mat AI machine learning approach

To aid in scoring the tangle maturity levels, we developed a “pathomics” approach using machine learning to recognize all tangle maturity levels. The Tangle-mat AI model achieved a macro-recall of 54% (**Supplementary Fig. 6a, Supplementary Fig. 7**). Most classification errors occurred between adjacent tangle maturity levels (i.e. pretangles with intermediary 1s, mature tangles with intermediary 1s and intermediary 2s, etc.). When adjacent classes were considered acceptable predictions, the macro-recall increased to 71% across maturity levels. Model macro-precision averaged 59% and increased to 78% when including adjacent classes (**Supplementary Fig. 6b**). When model-predicted tangle counts were converted to the semi-quantitative scale, results were similar to manual scoring (**Supplementary Fig. 6c**). Most of the differences between the manual tangle maturity scores and the Tangle-mat AI model were in adjacent scores. Using these tangle maturity scores, we then converted to the tangle maturity scale as described earlier (**Table 3**). When the model’s semi-quantitative scores were used to generate a tangle maturity scale, we found good agreement with the manually derived scale (**Supplementary Fig. 6d**). Analysis of the Tangle-mat AI model-derived tangle maturity scale revealed similar patterns to those observed using the manually derived scale (**Fig. 5, Supplementary Fig. 3c, 4c, 5c**). Specifically, the Tangle-mat AI maturity scale had moderate to strong correlations for all antibodies to Braak stage except MN423 in the subiculum and CA2/3. The model also correlated moderately to strongly for antibodies through the tangle lifespan with subsector thickness and neuronal count. Neuronal counts remained non-significant with the model-derived tangle maturity scale in the CA4 (**Supplementary Fig. 5c**).

Given the difficulty in delineating tangle maturity levels, a domain expert relabeled a subset of images to assess intra-rater performance compared to the model. Similar patterns emerged, with most confusion occurring between adjacent maturity levels. The intra-rater macro-recall was 64% across maturity levels, rising to 86% when including adjacent classes (**Supplementary Fig. 6e**). The intra-rater macro-precision was 63% and increased to 84% when including adjacent classes (**Supplementary Fig. 6f**).

### Tau burden correlations

To compare the tangle maturity quantification with more widely used methods, we performed a total tau pathology burden analysis using digital pathology to analyze annotated hippocampal subsectors. Quantification of the total burden of tau staining revealed a dramatic increase in phosphorylated tau antibodies and TOC1 as Braak stage increases compared to the other antibodies in the CA1, starting from Braak III to IV (**Fig. 7a**). Similar patterns were observed in the subiculum, CA2/3, and CA4 (**Supplementary Fig. 8a, 9a, 10a**).

As the total tau burden analysis did not distinguish between tangle and neuritic burden, we employed pattern recognition software (i.e., GENIE) to aid in this endeavor. For all hippocampal subsectors, the tangle burden remained below 10% (**Fig. 7, Supplementary Fig. 8b, 9b, 10b)**. Neuritic pathology accounted for a substantial increase in the burden of phosphorylated tau antibodies and TOC1 across Braak stages (**Fig. 7, Supplementary Fig. 8c, 9c, 10c**). Notably, pT181 burden was elevated in in the CA2/3 at Braak stage II compared to Braak stage III (**Supplementary Fig. 9a, c**), possibly reflecting physiologic axonal tau staining [50].

**Fig. 7.**
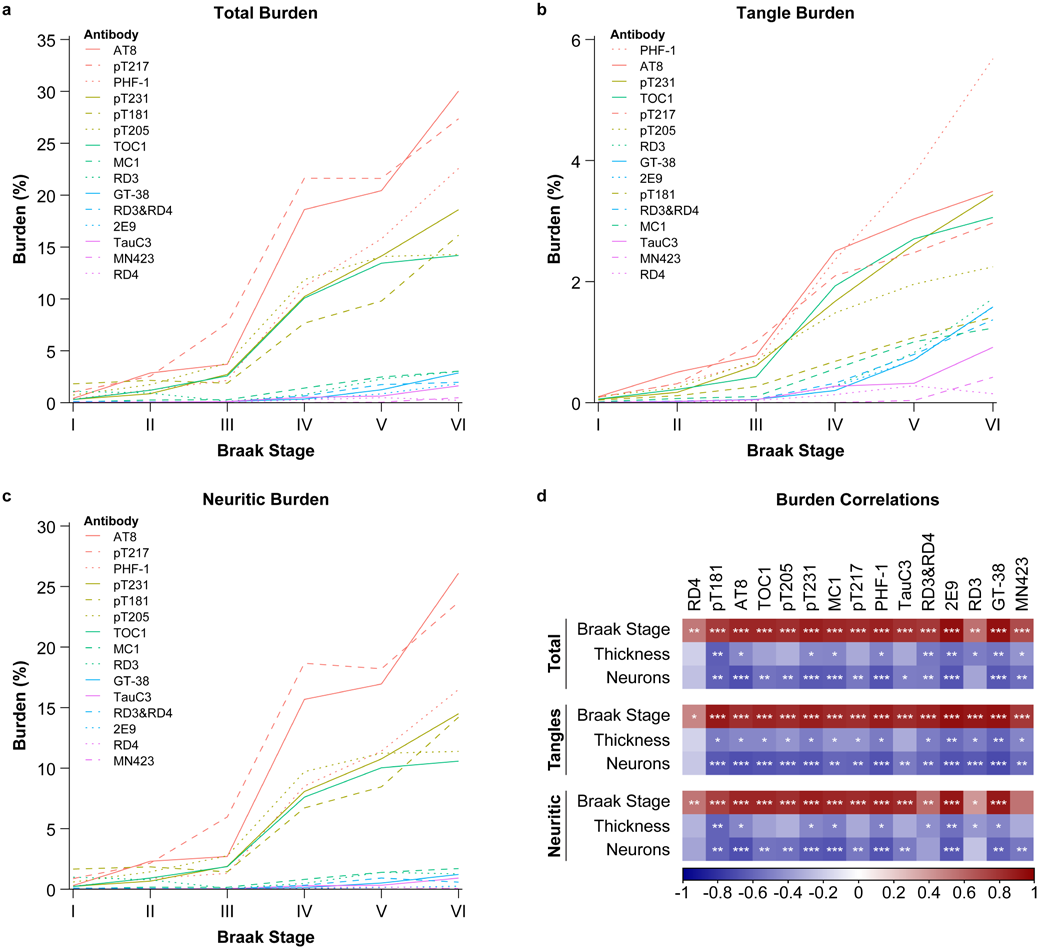
Pathomics-based tau burden analyses and quantification in the CA1. Pattern recognition software (i.e., GENIE) was used to subclassify tangle and neuritic burger from total tau immunoreactivity. Total, tangle, and neuritic burden in the CA1. Line graphs of (**a**) total burden, (**b**) tangle burden, and (**c**) neuritic burden in CA1 for each Braak stage. (**d**) Correlogram of Spearman correlations between total burden, tangle burden, or neuritic burden in the CA1 for each stain with neuronal count, CA1 thickness, and Braak Stage. The line graph key is organized by highest to lowest burden at Braak stage VI. *, p<0.05; **, p<0.01; ***, p<0.001.

In the CA1, Spearman correlations between total burden and Braak stage showed moderate to very strong correlations for all antibodies, with RD4 and RD3 having the weakest correlations (**Fig. 7d**). Similar patterns were observed in the other subsectors (**Supplementary Fig. 8d, 9d, 10d**). Tangle burden for all subsectors had moderate to very strong correlations with Braak stage (**Fig. 7, Supplementary Fig. 8d, 9d, 10d**). Neuritic burden for all antibodies had moderate to very strong correlations with Braak stage (**Fig. 7, Supplementary Fig. 8d, 9d, 10d**).

Total burden of AT8, pT181, pT231, MC1, PHF-1, RD3&RD4, 2E9, RD3, and MN423 had moderate correlations with CA1 thickness (**Fig. 7d**). Tangle burden of all antibodies except RD4 and TauC3 had moderate inverse correlations with CA1 thickness. CA1 thickness had moderate correlations with pT181, pT231, MC1, PHF-1, RD3&RD4, 2E9, RD3, and GT-38 neuritic burden. Total burden of all antibodies had moderate inverse correlations with subiculum thickness (**Supplementary Fig. 8d**). Tangle burden for all antibodies except TauC3 had moderate inverse correlations with subiculum thickness (**Supplementary Fig. 8d**). Neuritic burden in the subiculum had moderate correlations with thickness for all antibodies except TOC1, pT205, pT217, and RD3 (**Supplementary Fig. 8d**). CA2/3 thickness had moderate inverse correlations with total burden of AT8, pT231, and MC1 (**Supplementary Fig. 9d**). CA2/3 thickness had moderate inverse correlations with pT181, pT231, MC1, and GT-38 tangle burden (**Supplementary Fig. 9d**). Neuritic burden of AT8, MC1, and 2E9 had moderate inverse correlations with thickness in the CA2/3 (**Supplementary Fig. 9d**). Due to the anatomy of CA4 lacking parallel borders as it is encircled by the dentate fascia, no thickness measurements are available.

Neuronal counts had moderate inverse correlations with total, tangle, and neuritic burden with most antibodies in the CA1 (**Fig. 7d**). In the subiculum, neuronal counts had moderate inverse correlations with total, tangle, and neuritic burden for most antibodies (**Supplementary Fig. 8d**). In the CA2/3, neuronal counts had moderate inverse correlations with AT8, pT231, 2E9, and GT-38 total burden (**Supplementary Fig. 9d**). Tangle burden in the CA2/3 had moderate inverse correlations to neuronal count with pT231, 2E9, and GT-38, and neuritic burden had moderate inverse correlations with AT8, pT205, pT231, and 2E9 (**Supplementary Fig. 9**). Neuronal counts were not correlated with any total, tangle, or neuritic burden for antibodies in the CA4 (**Supplementary Fig. 10d**).

## Discussion

To gain deeper insights into posttranslational modifications of tau across the hypothesized lifespan of tangle-bearing neurons, we developed a novel tangle maturity scale to quantify tangle maturity levels in the human hippocampus. Using the proportion of the tangle maturity levels, we ordered the antibodies from early to advanced tangle maturity markers to aid in antibody selection for clinicopathology studies focused on AD-relevant tau accumulation. Applying the tangle maturity scale, we showed increasing correlation strength with neuronal counts and subsector thickness as tangle maturity markers become more advanced. We further developed pathomics approach using machine learning to classify the tangle maturity scores and scale, which performed similarly to manual scoring. These methods extend our knowledge in AD by providing morphology-based quantification of tangle maturity with biologically meaningful context.

We comprehensively characterized the tangle maturity levels of 15 antibodies and used the semi-quantitative tangle maturity scores to stratify the antibodies into “early” (mostly pretangles and mature tangles), “middling” (mature tangles with pretangles and ghost tangles), and “advanced” (mostly ghost tangles and mature tangles) tangle maturity markers. Previous studies characterized tangle maturity levels recognized by a selected set of tau antibodies [7, 17, 33, 35, 70, 71], providing further motivation to expand our understanding to a larger series of tau antibodies. Our results highlight recognition of tangle-associated neuritic clusters by AT8 and PHF-1 in proximity to ghost tangles, and replicate findings of pT231 as an early tangle maturity marker [7]. A recent study characterized the immunophenotype of tangles using immunofluorescence of 4R tau, AT8, 3R tau, and MN423 [35]. While our study did not include multiplex imaging, we observed similar results: 4R and AT8 tau staining pretangles, a mix of 4R, AT8, 3R tau, and MN423 in mature tangles, and 3R tau and MN423 in ghost tangles. Additionally, we replicated the findings of the 4R to 3R shift from pretangles to ghost tangles [35, 70], which may be of particular interest in the biomarker field with the advent of 4R tau PET radioligands [31]. Our results also confirm prior studies demonstrating the early emergence of specific tau conformations, including oligomeric forms of tau and that are linked to potential mechanisms of toxicity [16, 17, 33, 41, 61, 69, 71]. While the current study focused solely on progression through Braak stages with selected typical Alzheimer’s disease cases at higher stages, we look forward to continued progress in understanding tangle maturity in the context of selective vulnerability in non-amnestic forms of Alzheimer’s disease [6, 9, 11, 25].

We ordered antibodies from early to advanced based on the proportion of maturity levels observed for each antibody in the CA1. We chose the CA1 as this is the hippocampal subsector reported to accumulate neurofibrillary tangles the earliest compared to the rest of the hippocampus [13]. The results follow the hypothesized order of Alz-50, TauC3, Tau-66, and MN423 [10], but diverged for some based on previously reported hypothesized orders of the early tangle changes (pT231 & pS396/pS404 ➔ pS202/pT205 ➔ MC1 ➔ argyrophilic fibrils [4]; or pT231 ➔ Tau-C3 ➔ pT231 ➔ pS202/pT205 ➔ pT212/pS214 ➔ Alz50 ➔ tangle elongation [46]). Methodologic approaches may explain differences in ordering, as discrete assessment of regional tau positivity or as early molecular event may have different biological frequencies than proportion of tangle maturity levels shifting from early to advanced ordering. This may also suggest a partial temporal uncoupling of early tau posttranslational modifications during tangle development and progression, which highlights the need for integrated neuropathologic evaluation in longitudinal biomarker studies to define a more precise molecular chronology of tau pathogenesis.

Braak stage remains the most widely used framework for staging tau pathology in Alzheimer’s disease [12, 13], reflecting the topographic distribution of neurofibrillary tangles. To complement this approach with a morphology-based metric, we developed a semi-quantitative scoring system to assess the presence of each tangle maturity level and introduced the novel tangle maturity scale to summarize the relative predominance of tangle maturity levels. This framework enabled us to examine neurofibrillary tangle maturity as a variable, rather than aggregating all tau-positive pathology into a single burden measure. Using the tangle maturity scale, we identified consistent relationships between advancing tangle maturity markers and Braak stage, subsector thickness, and neuronal loss, particularly with increased ghost tangles associated with decreased thickness and fewer neurons in postmortem hippocampus. This agrees with previous studies demonstrating an association between ghost tangles and neuronal degeneration [18, 23]. Based on these collective observations, we recommend when possible that neuropathology-based studies consider using at least one early tangle maturity marker and one advanced tangle maturity marker. AT8, a widely used commercially available antibody, may facilitate greater cross-study comparisons on early tangle maturity, however for subtyping analysis aligned with advanced tangle maturity AT8 may not be the most effective choice for hippocampus [43]. For an advanced tangle maturity marker, we recommend commercially available 2E9 as mature tangles and ghost tangles are readily recognized. GT-38 may also be an effective advanced marker with undefined conformational shown to be specific for AD, but with a subtle note of caution that endstage ghost tangles may not be readily recognized [28, 43]. Selection of tangle maturity markers may be informed by general trend of stronger association between advanced tangle maturity markers and neuronal loss, or the relative preservation of tissue thickness observed with early 4R and select phosphorylated tau antibodies.

To support tangle maturity scoring using a computerized quantitative approach, we developed a machine learning model capable of recognizing the full spectrum of neurofibrillary tangle maturity levels (i.e., Tangle-mat AI). While prior machine learning models have successfully identified tangles using AT8 from hippocampus and prefrontal cortex with a total recall of 91% and precision of 72% on the testing cases [66]; differentiating tangle maturity levels was not the focus. A published model using primarily CP13, AT8, and PHF-1, was found to distinguish between pretangles and mature tangles but not ghost tangles despite successfully predicting Braak stage [72]. In the current study, we trained the model on 18 stains and recognized tangles using the 15 antibodies actively studied here to generalize the model across the tangle lifespan.

Tangle-mat AI scale values were moderately to strongly correlated with Braak stage for markers throughout the tangle lifespan. Additionally, the model performance paralleled intrarater reliability. The intra-rater reliability and class confusion between adjacent classes highlight the somewhat subjective nature of classifying tangles by maturity levels. This model may improve reproducibility of tangle quantification across studies. We observed similar correlations of the manual and model-derived tangle maturity scale with Braak stage, subsector thickness, and neuronal counts. Notably, MN423 showed fewer significant correlations, possibly due to the model’s difficulty detecting faintly stained ghost tangles, possibly confounded by signal from lipofuscin cross-reactivity in a subset of cases. Despite these challenges, the Tangle-mat AI model establishes a “pathomics” blueprint for reproducible, scalable, and morphologically informed tangle quantification that will accelerate digital pathology applications in Alzheimer’s research.

There are several strengths to our study, including a study series evenly distributed across Braak stages with sufficient tissue that enabled us to investigate across tangle maturity levels using 15 antibodies. Selective vulnerability of the hippocampus to tangle maturity was enhanced by subsector analysis of CA1, CA2/3, CA4, and subiculum, which are differentially affected by tau in Alzheimer’s disease (e.g., relative resistance of CA2/3). We performed a manual semi-quantitative approach in parallel with a pathomics approach applying machine learning and found similar performance for this complex endeavor. However, due to the cross-sectional nature of neuropathology studies, we were limited to examination of tangles at a single point in time. Mouse modeling of tauopathies may provide insights into early tangle progression that is more ideal for investigation of pretangles and mature tangles investigation [75, 77], whereas more advanced disease states may not replicate transcriptional and proteomic signatures found in human disease [60, 75]. To reduce variability, we solely focused on a single lamina (i.e., pyramidal layer) in an allocortical structure (i.e., posterior hippocampus), thus our findings need replication in mesocortical and isocortical regions to examine application of the tangle maturity scale. Future studies examining laminar distinction of tangle maturity levels may be greatly benefited by tissue clearing techniques [78]. Future refinement of the model, especially to account for staining artifacts and cellular context, will be essential for enhancing accuracy, supporting broader topographic application, and unlocking deeper insights into tau pathophysiology. Finally, it is important to consider that tangle maturity may not always progress in a monotonic fashion leading to death. Recent evidence from pS262 tau antibody investigation suggests a putative death state may be represented by presence of granulovacuolar degeneration in pretangles [38]. While we observed staining of tangle-bearing neurons occupied by GUDs, we did not observe immunoreactivity of granulovacuolar degeneration bodies as was reported with pS262 labelling [38]. Future work should consider inclusion of cell death markers to ascertain neuronal health state.

## Conclusions

This study introduces a novel framework for measuring neurofibrillary tangle maturity in Alzheimer’s disease, integrating semi-quantitative scoring with an AI-based model capable of recognizing morphologic tangle levels. By systematically classifying tau antibodies based on their maturity-level predilection, we provide knowledge toward marker selection that aligns with specific research goals (e.g., early detection, progression modeling). The tangle maturity scale, together with the Tangle-mat AI, enables nuanced, biologically informed analyses of tissue health. A transformative convergence of cross-disciplinary collaborations in neuropathology, digital pathology, AI & informatics, and biostatistics greatly accelerated progress toward our goal to integrate tissue-based measures with antemortem fluid and neuroimaging biomarkers in Alzheimer’s disease [55].

## Compliance with Ethical Standards

### Research involving Human Participants and/or Animals

All brains were acquired with appropriate ethical approval, and the study was approved by the Mayo Clinic institutional review board.

## Acknowledgements

We are grateful to the brain donors and their families for generous brain donations to help further our medical knowledge of Alzheimer’s disease and tau biology. We thank Monica Castanedes-Casey for histologic support and the Cytometry and Cell Imaging Core for imaging support. Programmatic support by Jessica Tranovich, Sabrina Rothberg, Avery Hatfield, and Dr. Kelsey Caetano-Anolles continues to be invaluable, and we are appreciative of their dedication. We thank Dr. Benjamin Wolozin for sharing the MC1 antibody. Research reported in this publication would not be possible without the dedicated support from the National Institute on Aging of the National Institutes of Health (R01-AG075802, R01-AG073282, RF1-AG069052, P30-AG062677), the National Institute of Neurological Disorders and Stroke of the National Institutes of Health (R01-NS082730), the Maibach Smiley Endowment, and the Alzheimer’s Association Florida Gulf Coast Chapter.

