## supplemental for "The Neurofibrillary Tangle Maturity Scale: A Novel Framework for Tangle Pathology Evaluation in Alzheimer’s Disease"

### Supplementary Information

Christina M. Moloney[1], Matthew H. Rutledge[2], Sydney A. Labuzan[1], Zhongwei Peng[2], Jessica F. Tranovich[1], Ashley C. Wood[1], Darren M. Rothberg[1], Ranjan Duara[3], Christian Lachner[4,5], Neill R. Graff-Radford[5], Dennis W. Dickson[1,6], Nicholas M. Kanaan[7], Rickey E. Carter[2#], Melissa E. Murray[1,6#]

[1] Department of Neuroscience, Mayo Clinic, Jacksonville, FL, 32224, USA
[2] Department of Quantitative Health Sciences, Mayo Clinic, Jacksonville, FL, 32224, USA
[3] Wien Center for Alzheimer's Disease and Memory Disorders, Mount Sinai Medical Center, Miami Beach, FL, USA
[4] Department of Psychiatry and Psychology, Mayo Clinic, Jacksonville, FL, 32224, USA
[5] Department of Neurology, Mayo Clinic, Jacksonville, FL, 32224, USA
[6] Department of Laboratory Medicine and Pathology, Mayo Clinic, Jacksonville, FL, 32224, USA
[7] Department of Translational Neuroscience, Michigan State University, MI, 49503, USA
[#] Corresponding authors

**Supplementary Table 1** Case demographics

| Case | Sex | Age | Braak Stage | Thal Phase | APOE ε4 | Clinical Diagnosis | Neuropathologic Diagnosis |
|---|---|---|---|---|---|---|---|
| 1 | F | 85 | I | 2 | ε4- | Normal | PART, possible |
| 2 | F | 92 | I | 0 | ε4- | Normal | PART, definite |
| 3 | M | 78 | I | 2 | ε4- | VaD | PART, possible/VaD |
| 4 | M | 88 | I | 1 | ε4- | VaD | PART, possible/VaD |
| 5 | F | 81 | II | 5 | ε4- | Depression | PA |
| 6 | F | 78 | II | 1 | ε4- | VaD | PART, possible/VaD |
| 7 | M | 83 | II | 4 | ε4- | AD/CVA | PA/VaD |
| 8 | M | 70 | II | 0 | ε4- | NPH | PART, definite |
| 9 | F | 87 | III | 3 | ε4- | Normal | PA |
| 10 | F | 89 | III | 3 | ε4- | Normal | SC/VaD/ARTAG |
| 11 | M | 88 | III | 0 | ε4- | MCI | PART, definite/VaD |
| 12 | M | 93 | III | 2 | ε4- | Normal | PART, possible |
| 13 | F | 82 | IV | 5 | ε4+ | AD | AD, early |
| 14 | F | 93 | IV | 5 | ε4- | DLB | AD, early/VaD |
| 15 | M | 84 | IV | 3 | ε4- | AD | AD, early/VaD |
| 16 | M | 83 | IV | 5 | ε4+ | AD | AD, early/VaD |
| 17 | F | 87 | V | 5 | ε4+ | AD | AD |
| 18 | F | 90 | V | 5 | ε4- | AD | AD |
| 19 | M | 82 | V | 5 | ε4+ | AD | AD/VaD |
| 20 | M | 84 | V | 4 | ε4+ | AD | AD/ VaD |
| 21 | F | 84 | VI | 5 | ε4- | AD | AD/VaD |
| 22 | F | 82 | VI | 5 | ε4- | AD | AD/VaD |
| 23 | M | 71 | VI | 5 | ε4+ | FTD/PPA | AD |
| 24 | M | 75 | VI | 5 | ε4+ | AD | AD |

Case characteristics in this series were reported in [38]. Acronyms: AD, Alzheimer's disease; ARTAG, age-related tau astrogliopathy; CVA, cerebrovascular accident; DLB, dementia with Lewy bodies; FTD, frontotemporal dementia; MCI, mild cognitive impairment; PA, pathologic aging; PART, primary age related tauopathy; PPA, primary progressive aphasia; SC, senile change; VaD (clinical), vascular dementia; VaD (neuropath), vascular disease

**Supplementary Table 2** Color deconvolution macro specifications

| Antibody | 2E9 | AT8 | GT-38 | MC1 | MN423 | PHF-1 | RD3 | RD3&RD4 | RD4 | TauC3 | TOC1 |
|---|---|---|---|---|---|---|---|---|---|---|---|
| Version | 9.1 | 9.1 | 9.1 | 9.1 | 9.1 | 9.1 | 9.1 | 9.1 | 9.1 | 9.1 | 9.1 |
| View Width | 1000 | 1000 | 1000 | 1000 | 1000 | 1000 | 1000 | 1000 | 1000 | 1000 | 1000 |
| View Height | 1000 | 1000 | 1000 | 1000 | 1000 | 1000 | 1000 | 1000 | 1000 | 1000 | 1000 |
| Overlap Size | 0 | 0 | 0 | 0 | 0 | 0 | 0 | 0 | 0 | 0 | 0 |
| Image Zoom | 1 | 1 | 1 | 1 | 1 | 1 | 1 | 1 | 1 | 1 | 1 |
| Classifier Neighborhood | 0 | 0 | 0 | 0 | 0 | 0 | 0 | 0 | 0 | 0 | 0 |
| Pixel Area ($mm^2$) | 2.53E-07 | 2.53E-07 | 2.53E-07 | 2.53E-07 | 2.53E-07 | 2.53E-07 | 2.53E-07 | 2.53E-07 | 2.53E-07 | 2.53E-07 | 2.53E-07 |
| Color Channel to Quantitate | 3 | 3 | 3 | 3 | 3 | 3 | 3 | 3 | 3 | 3 | 3 |
| Markup Image Type | 0 | 0 | 0 | 0 | 0 | 0 | 0 | 0 | 0 | 0 | 0 |
| Weak Positive Threshold | 235 | 235 | 235 | 235 | 235 | 235 | 235 | 235 | 235 | 235 | 235 |
| Medium Positive Threshold | 110 | 180 | 185 | 180 | 160 | 180 | 180 | 160 | 160 | 180 | 180 |
| Strong Positive Threshold | 110 | 180 | 185 | 180 | 160 | 180 | 180 | 160 | 160 | 180 | 180 |
| Black Threshold | 0 | 0 | 0 | 0 | 0 | 0 | 0 | 0 | 0 | 0 | 0 |
| Color(1) Red OD | 0.65 | 0.555 | 0.65 | 0.65 | 0.65 | 0.65 | 0.65 | 0.65 | 0.65 | 0.65 | 0.65 |
| Color(1) Green OD | 0.704 | 0.615 | 0.704 | 0.704 | 0.704 | 0.704 | 0.704 | 0.704 | 0.704 | 0.704 | 0.704 |
| Color(1) Blue OD | 0.286 | 0.561 | 0.286 | 0.286 | 0.286 | 0.286 | 0.286 | 0.286 | 0.286 | 0.286 | 0.286 |
| Color(2) Red OD | 7.20E-02 | 7.20E-02 | 7.20E-02 | 7.20E-02 | 7.20E-02 | 7.20E-02 | 7.20E-02 | 7.20E-02 | 7.20E-02 | 7.20E-02 | 7.20E-02 |
| Color(2) Green OD | 0.99 | 0.99 | 0.99 | 0.99 | 0.99 | 0.99 | 0.99 | 0.99 | 0.99 | 0.99 | 0.99 |
| Color(2) Blue OD | 0.105 | 0.105 | 0.105 | 0.105 | 0.105 | 0.105 | 0.105 | 0.105 | 0.105 | 0.105 | 0.105 |
| Color(3) Red OD | 0.417 | 0.268 | 0.341 | 0.341 | 0.341 | 0.341 | 0.341 | 0.341 | 0.341 | 0.341 | 0.341 |
| Color(3) Green OD | 0.572 | 0.57 | 0.575 | 0.575 | 0.575 | 0.575 | 0.575 | 0.575 | 0.575 | 0.575 | 0.575 |
| Color(3) Blue OD | 0.707 | 0.776 | 0.743 | 0.743 | 0.743 | 0.743 | 0.743 | 0.743 | 0.743 | 0.743 | 0.743 |

**Supplementary Table 3** Positive pixel count macro specifications

| Antibody | pT181 | pT205 | pT217 | pT231 |
|---|---|---|---|---|
| Version | 9.1 | 9.1 | 9.1 | 9.1 |
| View Width | 1000 | 1000 | 1000 | 1000 |
| View Height | 1000 | 1000 | 1000 | 1000 |
| Overlap Size | 0 | 0 | 0 | 0 |
| Image Zoom | 1 | 1 | 1 | 1 |
| Classifier Neighborhood | 0 | 0 | 0 | 0 |
| Pixel Area (millimeter-squared) | 2.53E-07 | 2.53E-07 | 2.53E-07 | 2.53E-07 |
| Hue Value (Center) | 0.1 | 0.1 | 0.1 | 0.1 |
| Hue Width | 0.1 | 0.1 | 0.1 | 0.1 |
| Color Saturation Threshold | 8.00E-02 | 8.00E-02 | 8.00E-02 | 8.00E-02 |
| Intensity Threshold WEAK (Upper Limit) | 235 | 235 | 235 | 235 |
| Intensity Threshold WEAK (Lower Limit) | 235 | 235 | 235 | 235 |
| Intensity Threshold MEDIUM (Upper Limit) | 235 | 235 | 235 | 235 |
| Intensity Threshold MEDIUM (Lower Limit) | 235 | 235 | 235 | 235 |
| Intensity Threshold STRONG (Upper Limit) | 235 | 235 | 235 | 235 |
| Intensity Threshold STRONG (Lower Limit) | 0 | 0 | 0 | 0 |
| Intensity Threshold Negative Pixels | -1 | -1 | -1 | -1 |

**Supplementary Table 4** Tangle-mat AI training antibodies.

| **Antibody** | **Vendor** | **Catalog #** | **Count** |
|---|---|---|---|
| AT8 | Thermo Fisher | MN1020 | 769 |
| 2E9 | Novus Biologicals | NBP2-25162 | 741 |
| PHF-1 | Peter Davies | N/A | 629 |
| RD3 | Millipore | 05-803 | 103 |
| GT-38 | Abcam | ab246808 | 97 |
| AT270 | Thermo Fisher | MN1050 | 81 |
| pT217 | Thermo Fisher | 44-744 | 63 |
| RD4 | Millipore | 05-804 | 56 |
| SERPINA5 | R&D | MAB1266 | 54 |
| MN423 | Michael Novak | | 50 |
| pT205 | Thermo Fisher | 44-738G | 40 |
| MC1 | Benjamin Wolozin | | 38 |
| AT180 | Thermo Fisher | MN1040 | 35 |
| TauC3 | Nicholas Kanaan | | 34 |
| RD3&RD4 | Millipore | 05-803&05-804 | 15 |
| Tau-66 | Nicholas Kanaan | | 9 |
| GT-38 | Virginia Lee | | 8 |
| 4R Tau | Cosmobio | CAC-TIP-4RT-P01 | 5 |

**Supplementary Table 5** Number of objects trained in Tangle-mat AI.

| **Class** | **Number of Objects** |
|---|---|
| Pretangle | 1406 |
| Intermediary 1 | 1860 |
| Mature | 2636 |
| Intermediary 2 | 695 |
| Ghost | 634 |
| Unclassified | 3682 |

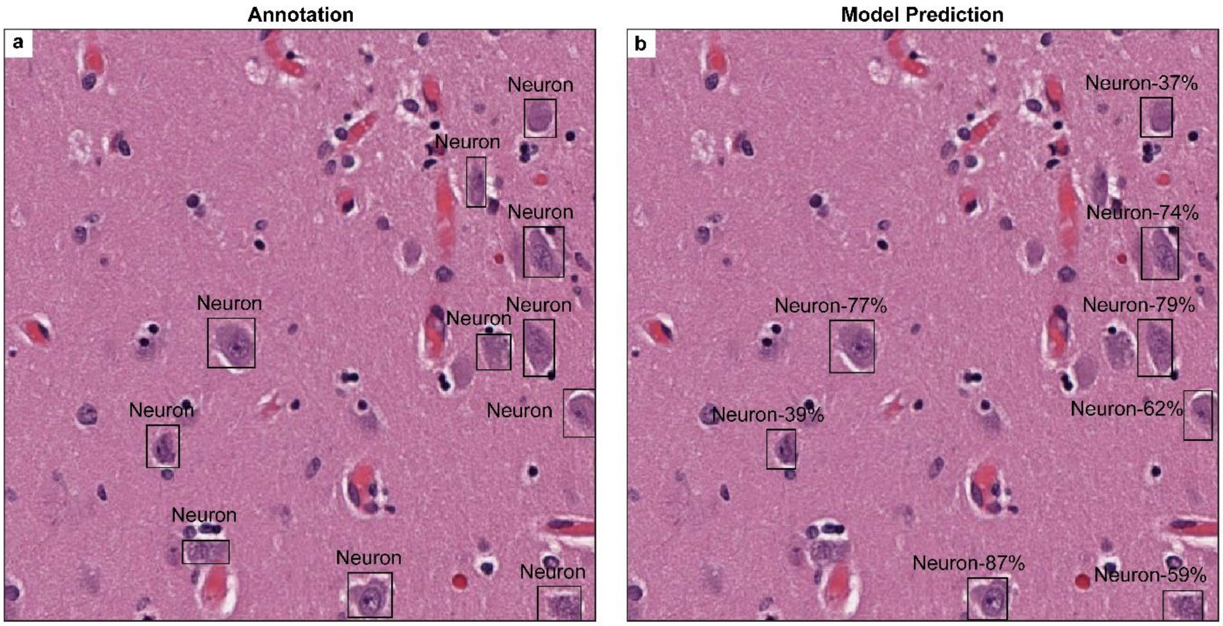

**Supplementary Fig. 1** Neuron-AI predictions. Example tiles of the neuron model with (**a**) annotations and (**b**) model predictions in hematoxylin and eosin-stained diagnostic slides.

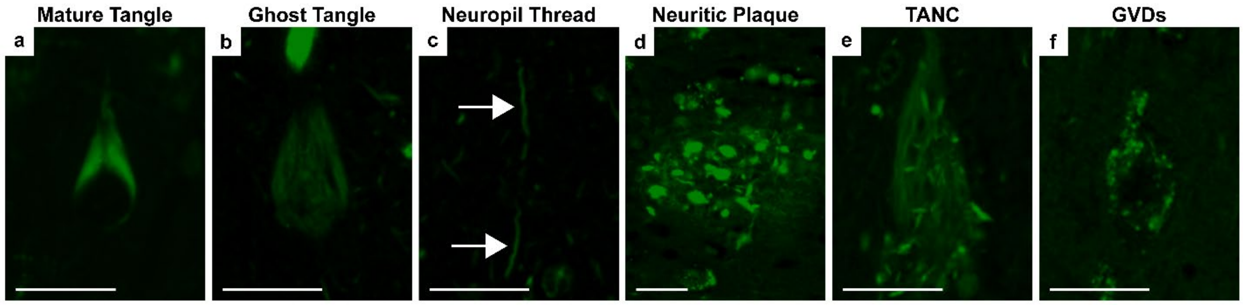

**Supplementary Fig. 2** Thioflavin-S tangle and non-tangle pathology. Representative images of a thioflavin-S positive (**a**) mature tangle, (**b**) ghost tangle, (**c**) neuropil thread (arrow), (**d**) neuritic plaque, (**e**) tangle-associated neuritic cluster, and (f) tangle-bearing neuron with granulovacuolar degeneration. No pretangles were observed. Scale bar measures 25 µm. Acronyms: TANC, tangle-associated neuritic cluster; GVD, granulovacuolar degeneration.

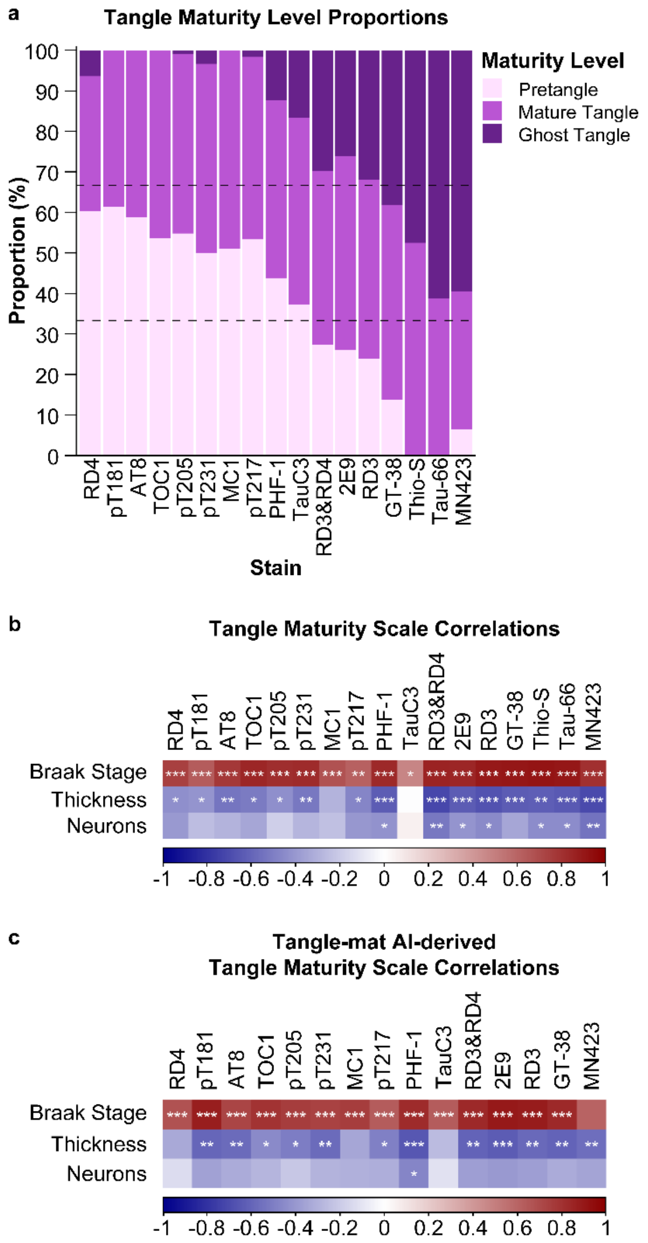

**Supplementary Fig. 3** Tangle maturity quantification and correlations in the subiculum. (**a**) Stacked bar graph of the proportion of pretangles, mature tangles, and ghost tangles for each stain. (**b**) Correlogram of Spearman correlations of the tangle maturity scale in the subiculum with Braak stage and tissue health (i.e., subiculum thickness, neurons) (**Fig. 5**). (**c**) Correlogram of Spearman correlations using the model derived tangle maturity scale. *, p<0.05; **, p<0.01; ***, p<0.001.

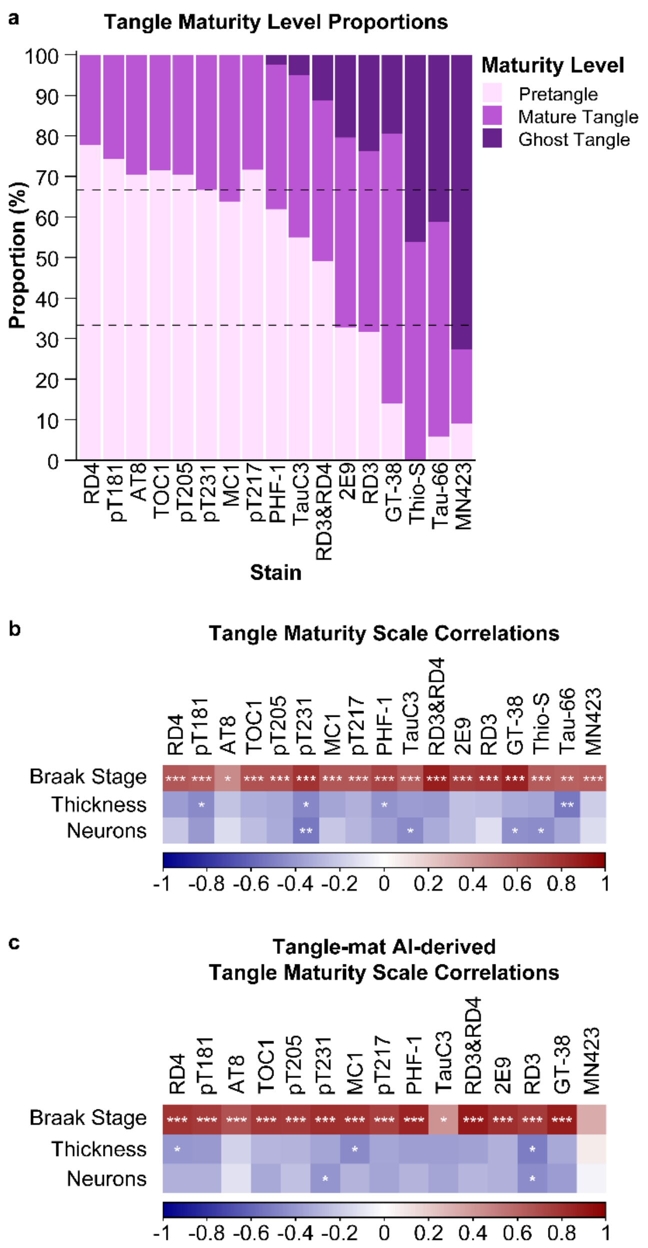

**Supplementary Fig. 4** Tangle maturity quantification and correlations in the CA2/3. (**a**) Stacked bar graph of the proportion of pretangles, mature tangles, and ghost tangles for each stain. (**b**) Correlogram of Spearman correlations of the tangle maturity scale in the CA2/3 with Braak stage and tissue health (i.e., CA2/3 thickness, neurons). All plots were ordered based on the order of stains in the CA1 (**Fig. 5**). (**c**) Correlogram of Spearman correlations using the model derived tangle maturity scale. *, p<0.05; **, p<0.01; ***, p<0.001.

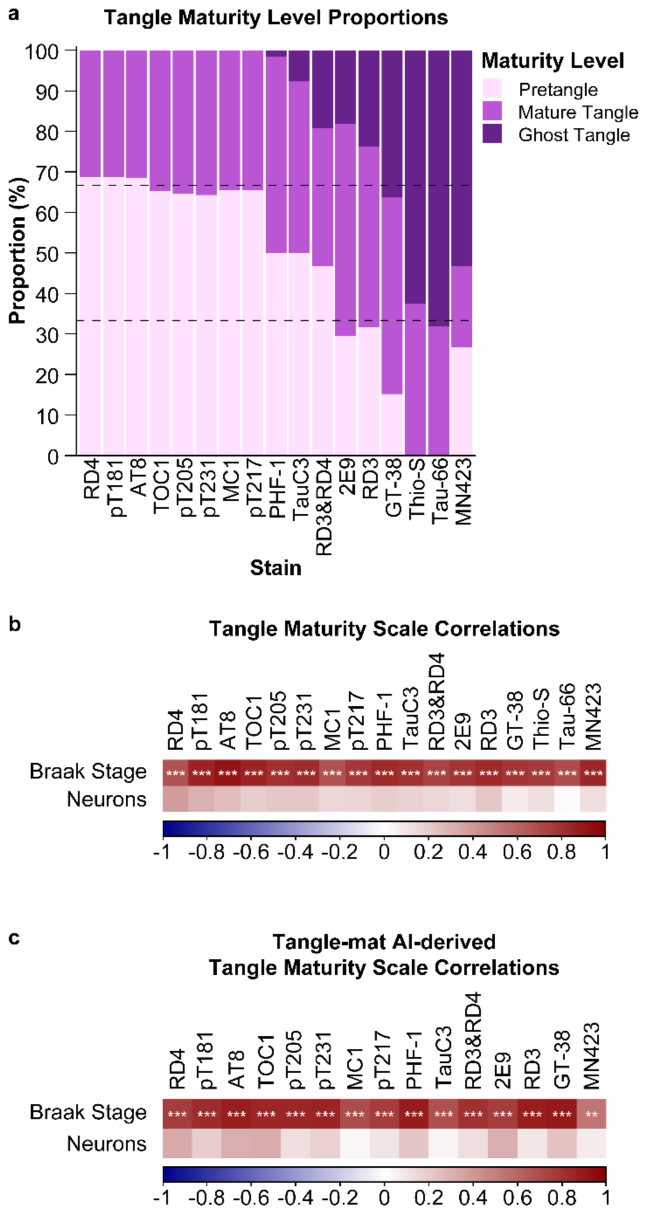

**Supplementary Fig. 5** Tangle maturity quantification and correlations in the CA4. (**a**) Stacked bar graph of the proportion of pretangles, mature tangles, and ghost tangles for each stain. (**b**) Correlogram of Spearman correlations of the tangle maturity scale in the CA4 with Braak stage and tissue health (i.e., neurons). All plots were ordered based on the order of stains in the CA1 (**Fig. 5**). (**c**) Correlogram of Spearman correlations using the model derived tangle maturity scale. *, $p<0.05$; **, $p<0.01$; ***, $p<0.001$.

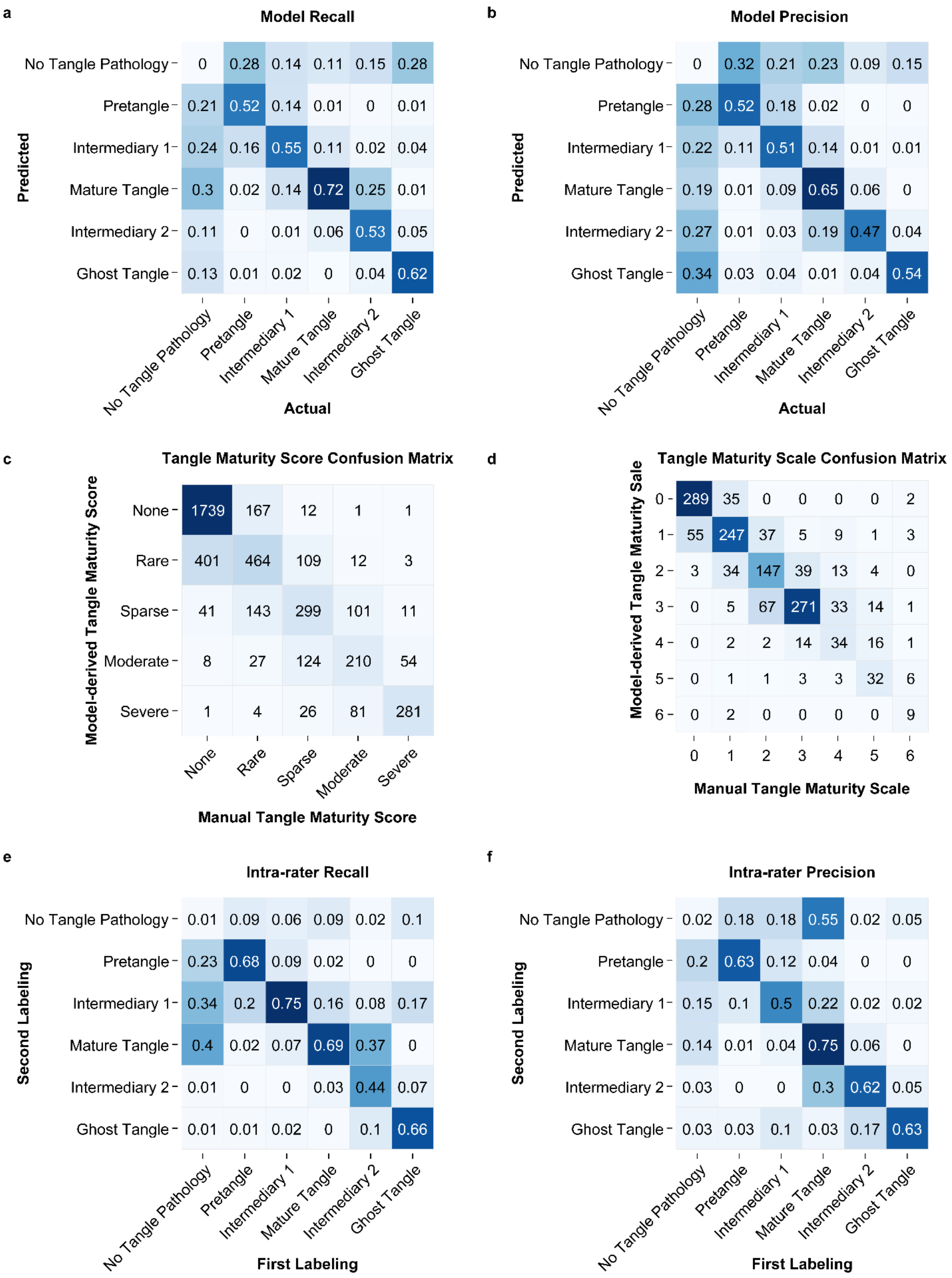

**Supplementary Fig. 6** Confusion matrices showing (**a**) model recall for 5-fold cross validation, (**b**) model precision for 5-fold cross validation, (**c**) human derived semi-quantitative scores and model generated semi-quantitative scores, and (**d**) human derived tangle maturity scale and model generated tangle maturity scale. Confusion matrices comparing (**e**) intra-rater recall, as well as (**f**) intra-rater precision. The strength of the correlation is noted with increasing intensity of blue in the cell. Overall, there was close agreement with most differences in adjacent classes. There were a few outliers where the scales were quite different, which was more commonly observed in advanced tangle maturity markers. Cells from each fold were summed to create the final table and cases where the model correctly predicted an empty box were omitted for clarity.

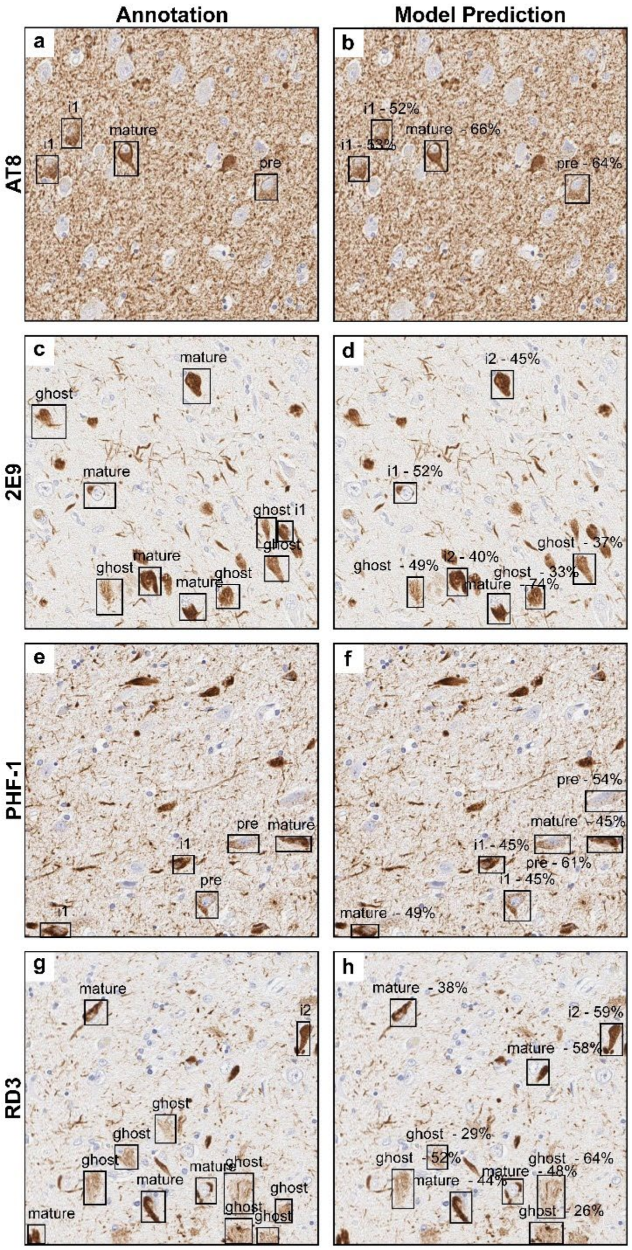

**Supplementary Fig. 7** Example tiles of the Tangle-mat AI model. (**a**) Manual annotation identified by black rectangle and (**b**) model prediction of tangles maturity level accompanied by and of an AT8 tile. (**c**) Annotation and (**d**) model prediction of a 2E9 tile. (**e**) Annotation and (**f**) model prediction of a PHF-1 tile. (**g**) Annotation and (**h**) model prediction of a RD3 tile. Abbreviations: pre, pretangle; i1, intermediary 1; i2, intermediary 2; Tangle-mat AI, tangle maturity artificial intelligence model.

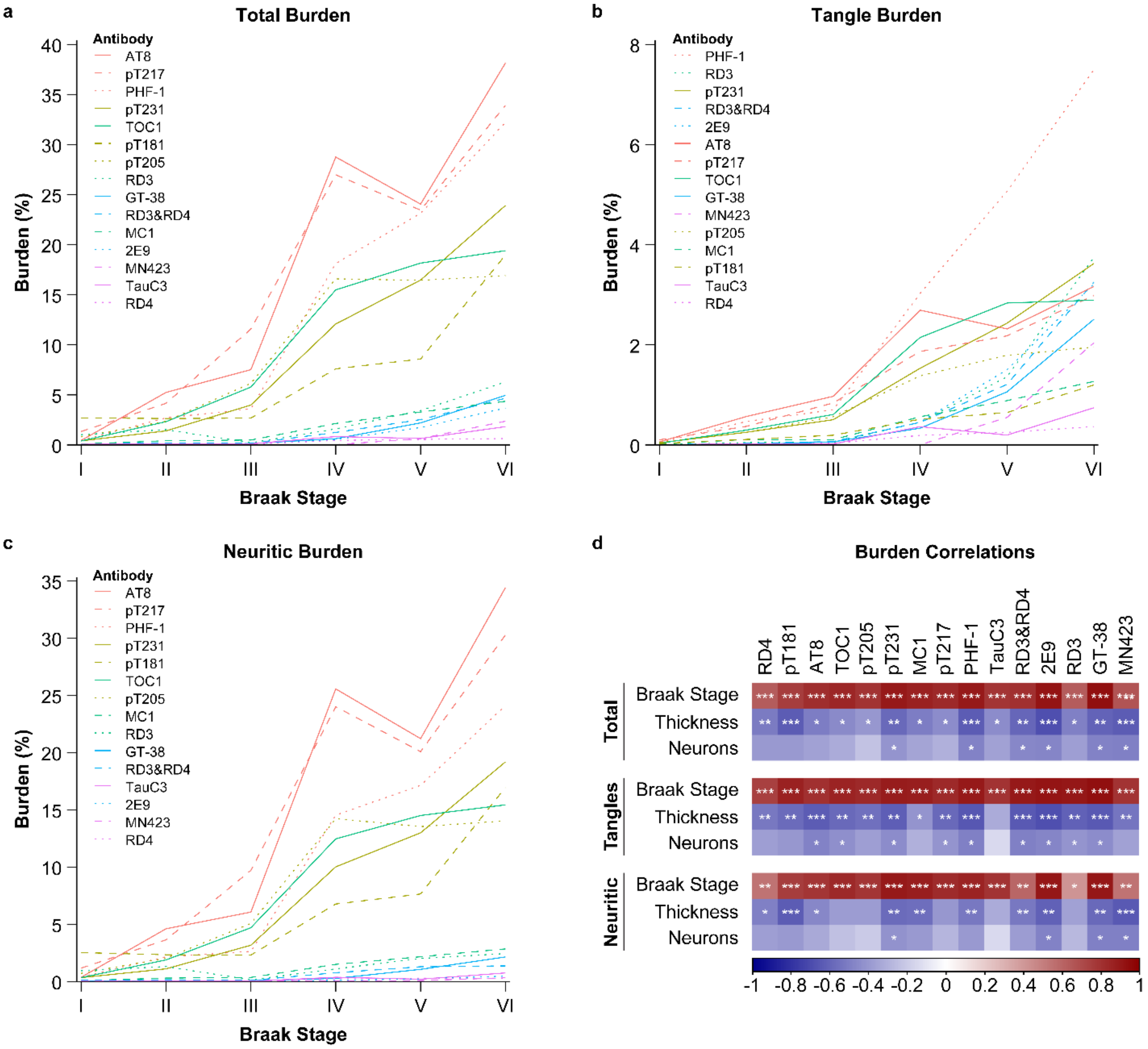

**Supplementary Fig. 8** Burden analyses and quantification in the subiculum. Total, tangle, and neuritic burden in the subiculum. Line graph of (**a**) total burden, (**b**) tangle burden, and (**c**) neuritic burden in subiculum for each Braak stage. (**d**) Correlogram of Spearman correlations between total burden, tangle burden, or neuritic burden in the subiculum for each stain with neuronal count, subiculum thickness, and Braak Stage. The line graph key is organized by highest to lowest burden at Braak stage VI. *, p<0.05; **, p<0.01; ***, p<0.001.

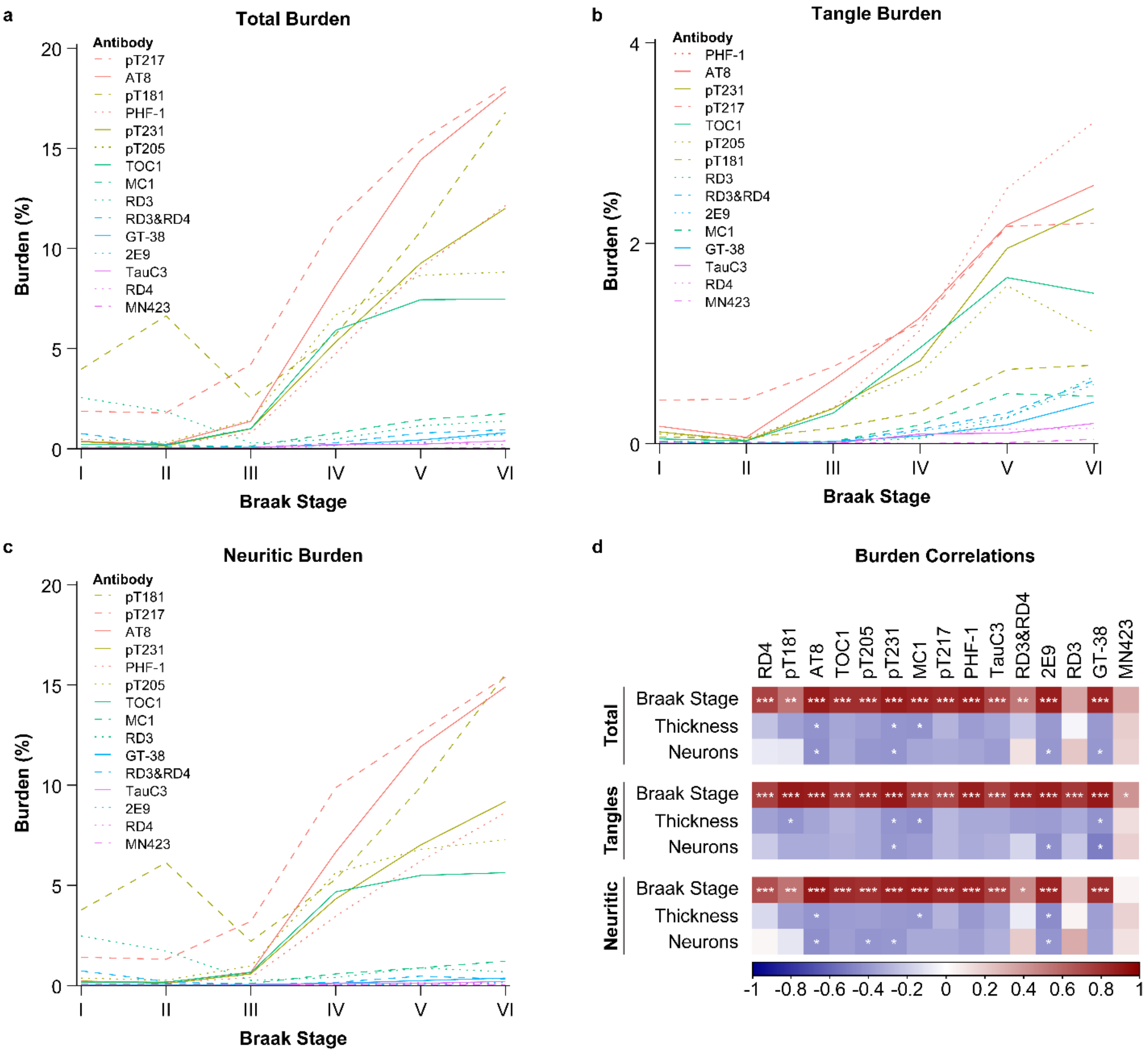

**Supplementary Fig. 9** Burden analyses and quantification in the CA2/3. Total, tangle, and neuritic burden in the CA2/3. Line graph of (**a**) total burden, (**b**) tangle burden, and (**c**) neuritic burden in CA2/3 for each Braak stage. (**d**) Correlogram of Spearman correlations between total burden, tangle burden, or neuritic burden in the CA2/3 for each stain with neuronal count, CA2/3 thickness, and Braak Stage. The line graph key is organized by highest to lowest burden at Braak stage VI. *, $p<0.05$; **, $p<0.01$; ***, $p<0.001$.

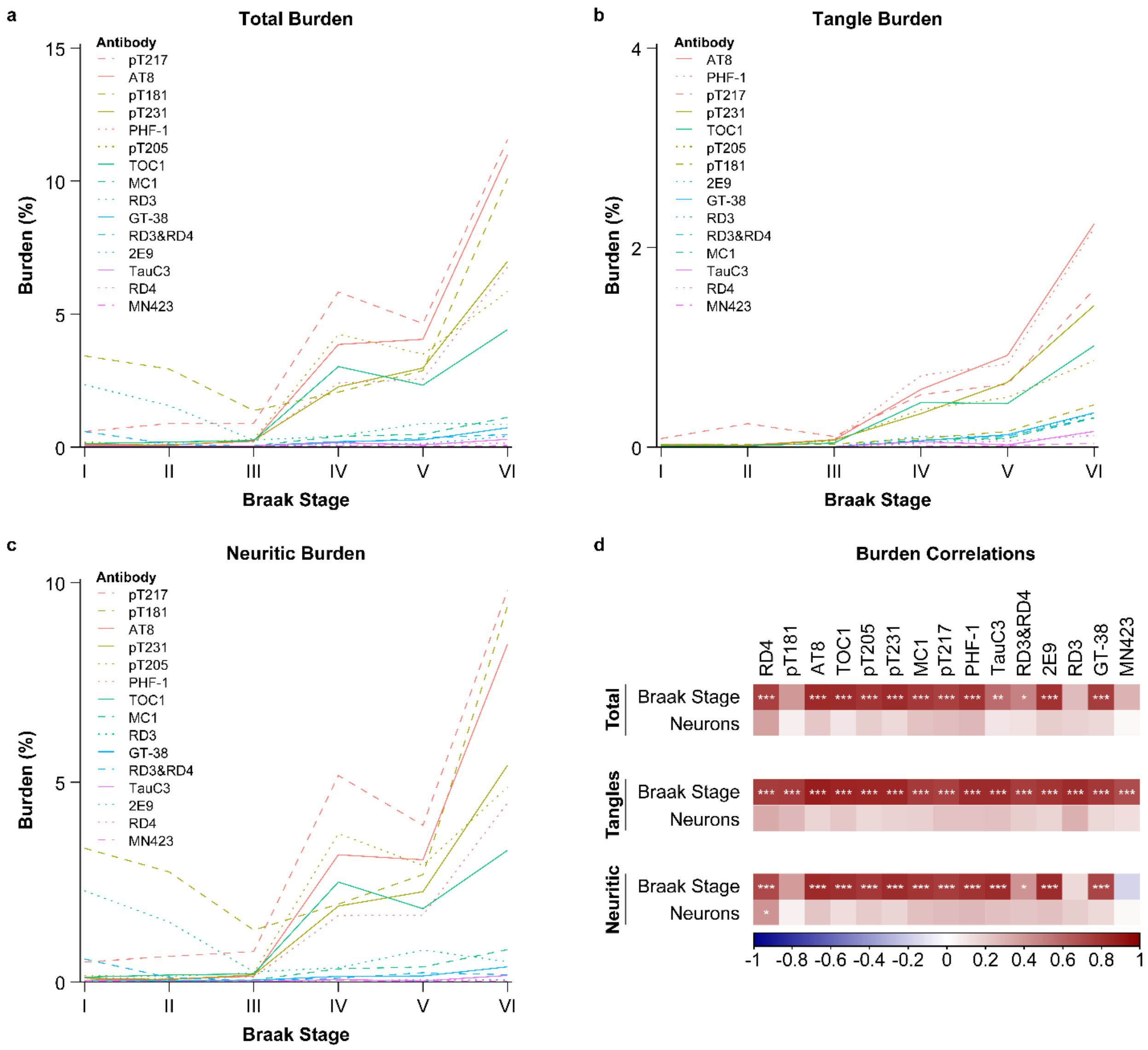

**Supplementary Fig. 10** Burden analyses and quantification in the CA4. Total, tangle, and neuritic burden in the CA4. Line graph of (**a**) total burden, (**b**) tangle burden, and (**c**) neuritic burden in CA4 for each Braak stage. (**d**) Correlogram of Spearman correlations between total burden, tangle burden, or neuritic burden in the CA4 for each stain with neuronal count and Braak stage. The line graph key is organized by highest to lowest burden at Braak stage VI. *, p<0.05; **, p<0.01; ***, p<0.001.
